# Copy number-aware deconvolution of tumor-normal DNA methylation profiles

**DOI:** 10.1101/2020.11.03.366252

**Authors:** Elizabeth Larose Cadieux, Nana E. Mensah, Carla Castignani, Miljana Tanić, Gareth A. Wilson, Michelle Dietzen, Pawan Dhami, Heli Vaikkinen, Annelien Verfaillie, Cristina Cotobal Martin, Toby Baker, Thomas B. K. Watkins, Selvaraju Veeriah, Mariam Jamal-Hanjani, Nnennaya Kanu, Nicholas McGranahan, Andrew Feber, TRACERx Consortium, Charles Swanton, Stephan Beck, Jonas Demeulemeester, Peter Van Loo

## Abstract

Aberrant methylation is a hallmark of cancer, but bulk tumor data is confounded by admixed normal cells and copy number changes. Here, we introduce Copy number-Aware Methylation Deconvolution Analysis of Cancers (CAMDAC; https://github.com/VanLoo-lab/CAMDAC), which outputs tumor purity, allele-specific copy number and deconvolved methylation estimates. We apply CAMDAC to 122 multi-region samples from 38 TRACERx non-small cell lung cancers profiled by reduced representation bisulfite sequencing. CAMDAC copy number profiles parallel those derived from genome sequencing and highlight widespread chromosomal instability. Deconvolved polymorphism-independent methylation rates enable unbiased tumor-normal and tumor-tumor differential methylation calling. Read-phasing validates CAMDAC methylation rates and directly links genotype and epitype. We show increased epigenetic instability in adenocarcinoma *vs.* squamous cell carcinoma, frequent hypermethylation at sites carrying somatic mutations, and parallel copy number losses and methylation changes at imprinted loci. Unlike bulk methylomes, CAMDAC profiles recapitulate tumor phylogenies and evidence distinct patterns of epigenetic heterogeneity in lung cancer.

## INTRODUCTION

Like DNA mutations, epigenetic alterations, such as disruption of DNA methylation and chromatin architecture, are now acknowledged as a universal feature of tumorigenesis (Baylin and Jones, 2016; Feinberg et al., 2016; Shen and Laird, 2013). These changes accumulate in normal cells throughout their lifetime (Hannum et al., 2013; Hao et al., 2016; Horvath, 2013) and while most have no effect, a small subset may provide a selective advantage for the cell (Feinberg et al., 2006; Flavahan et al., 2017). Through successive gains of hallmark cancer cellular capabilities (Hanahan and Weinberg, 2011), cells may ultimately acquire a fully malignant phenotype, while continuing to evolve in response to environmental pressures (Quail and Joyce, 2013).

DNA methylation is a mitotically heritable covalent DNA modification. In vertebrates, cytosines (C) can be methylated to form 5-methylcytosines (5mC), mostly in a CpG context (Bird, 2002; Smith and Meissner, 2013). This methylation occurs throughout the entire genome where it may aid to suppress cryptic transcription (Costello et al., 2010) and keep mobile genetic elements in check (Deniz et al., 2019). Active regulatory regions often contain unmethylated CpG-islands, irrespective of the expression level of associated genes (Deaton and Bird, 2011; Eckhardt et al., 2006; Ziller et al., 2013), although methylation of these CpG-rich features is usually anti-correlated with transcription factor binding. Genomic alterations removing binding sites enable DNA methyltransferase activity to spread and deplete gene expression (Smith and Meissner, 2013); *vice versa*, aberrations creating new binding sites can stimulate transcription (Huang et al., 2013; Mansour et al., 2014).

The cancer methylome displays characteristics from its cell of origin in addition to somatic DNA methylation changes (Greger et al., 1989; Wen et al., 2009). As methylation profiling provides insight into (disease) cell states and other aspects of biology, with a readout that is not reliant on live cells nor large amounts of high quality material, it may yield powerful biomarkers (Feber et al., 2017; Heyn and Esteller, 2012; Koch et al., 2018).

Massively parallel sequencing efforts by the Cancer Genome Atlas (TCGA) and the International Cancer Genome Consortium (ICGC) have revealed somatic alterations across thousands of cancer genomes (ICGC/TCGA Pan-Cancer Analysis of Whole Genomes Consortium, 2020; Hutter and Zenklusen, 2018). Although whole-genome (WGBS, Lister et al., 2009) and reduced-representation bisulfite sequencing (RRBS, Meissner, 2005) are gaining in popularity (Hansen et al., 2011; Hu et al., 2021; Pfister et al., 2014; Sun et al., 2015; Zhao et al., 2020; Ziller et al., 2013), the cancer methylome is considerably less well charted. Intra-tumor DNA methylation heterogeneity has been widely reported (Brocks et al., 2014; Hua et al., 2020; Klughammer et al., 2018; Mazor et al., 2015), however the interpretation of bulk tumor methylation data is hampered by the admixture of normal cells (Chakravarthy et al., 2018) and the occurrence of somatic copy number alterations (CNAs) in most cancers (Martin-Trujillo et al., 2017). While a number of methods have been developed to assess normal cell contributions (Barrett et al., 2017; Guo et al., 2017; Teschendorff et al., 2017; Zheng et al., 2014), research focus has generally been restricted to high purity samples and CNA-quiet cancer types such as Ewing sarcoma (Sheffield et al., 2017) and malignancies of the central nervous system (Capper et al., 2018) or lymphoid origins (Kulis et al., 2012; Landau et al., 2014; Nordlund et al., 2013; Oakes et al., 2016). Similarly, accurate estimates of purity and copy number are critical when querying intra-tumor heterogeneity and reconstructing tumor evolutionary histories (Tarabichi et al., 2021). Current metrics to quantify tumor heterogeneity from bisulfite sequencing data also assume high tumor content and/or the absence of CNAs (Chen et al., 2021; Landan et al., 2012; Landau et al., 2014; Li et al., 2016; Sheffield et al., 2017).

To address these issues, we developed a tool for Copy number-Aware Methylation Deconvolution Analysis of Cancers (CAMDAC). CAMDAC provides accurate allele-specific copy number and purity estimates from tumor RRBS data. Formalizing the relationship between methylation rates, copy number and tumor purity, CAMDAC extracts purified tumor methylomes from bulk tumor and tissue-matched normal bisulfite sequencing data. The corrected tumor methylation rates allow for accurate quantification of differential methylation, both between tumor and normal cells and between different tumors or sampled regions. Phasing to sequence variants, we assess epigenetic instability and the interplay between DNA methylation and somatic copy number and single-nucleotide variants. CAMDAC deconvolved (allele-specific) tumor methylation profiles reveal intra-tumor subclonal relationships and allow to quantify allele-specific methylation signals.

## RESULTS

### Bulk tumor methylation rates are confounded by tumor purity and copy number

To study epigenetic inter- and intra-tumor heterogeneity in non-small cell lung cancer, we obtained surgically resected primary tumor samples from 24 lung adenocarcinoma (LUAD) and 14 squamous cell lung carcinoma (LUSC) patients (**Table S1**) from the TRACERx 100 cohort, and performed multi-region RRBS of 122 tumor regions in total (2-7 per patient, **Table S2**). As a normal reference, we included samples of adjacent normal tissue for 37/38 patients.

We hypothesize that normal cell admixture (tumor purity) and somatic copy number alterations affect the methylation rate of bulk tumor samples (**Figure 1A**). We set out to assess this effect knowing that previous work using whole-exome sequencing (WES) of the same samples revealed considerable variability in purity and copy number (Jamal-Hanjani et al., 2017). We selected CpG loci that are confidently unmethylated in the adjacent normal sample (posterior 99% highest density interval on methylation rate HDI^99^ ⊆ [0, 0.2], **Methods**), stratified them by allele-specific copy number state in three tumor regions with different purity, and evaluated their bulk methylation rates (**Figure 1B**). The majority of these loci exhibit bulk methylation rates close to 0 (88% with HDI^99^ ⊆ [0, 0.2]), suggesting that most sites are not differentially methylated in the tumor. A second population of CpG sites is visible across all samples, the modal methylation rate of which shifts with tumor purity and copy number. We observed a similar effect evaluating CpG loci that are confidently methylated in the adjacent normal sample (**Figure S1**). These observations confirm that normal cell admixture and copy number changes confound tumor methylation signals. We therefore set out to infer tumor purity and copy number from bulk tumor RRBS data, and extract purified tumor methylation rates, unpolluted with signals from non-tumor cells.

**Figure 1.**
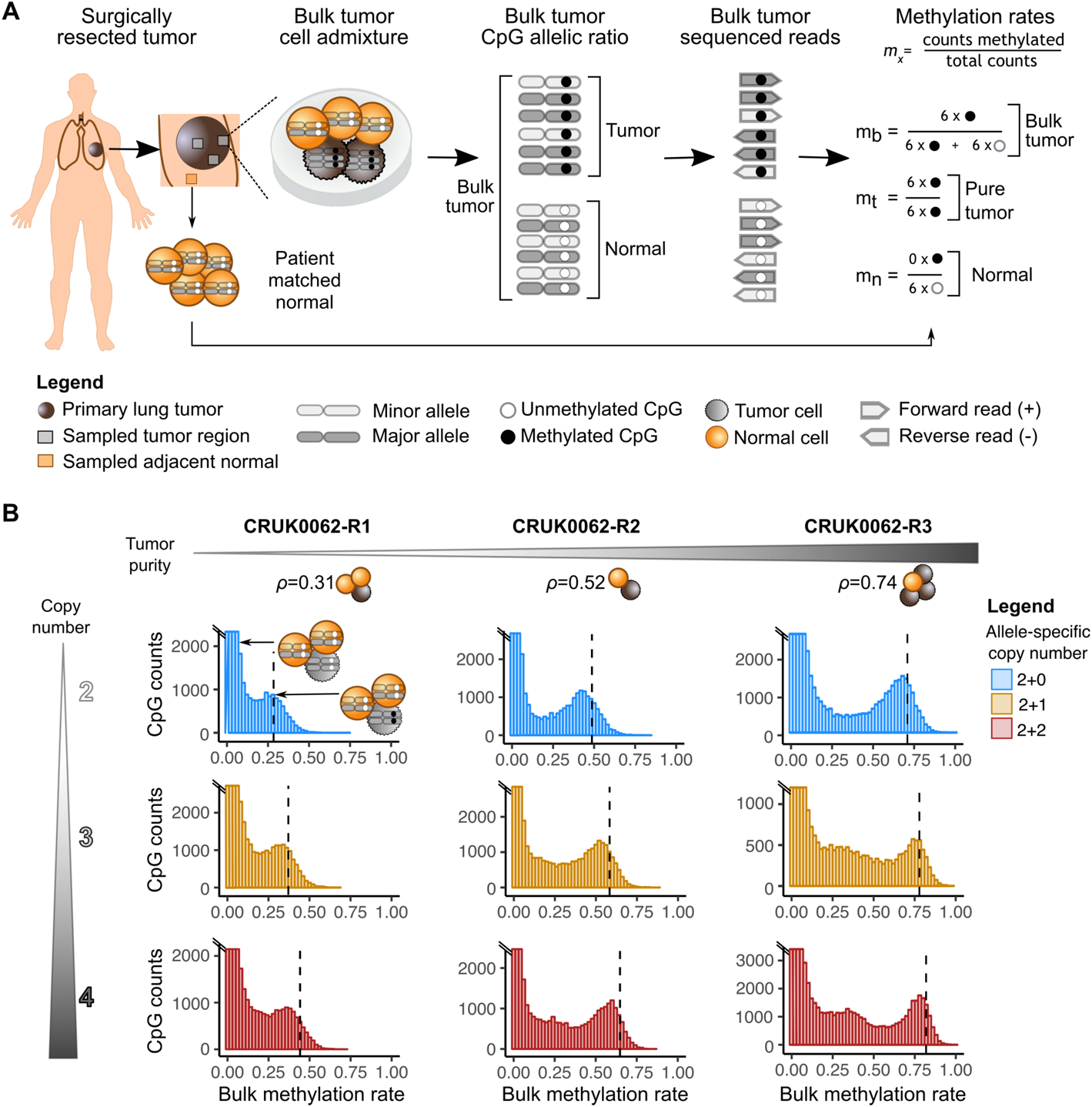
Tumor purity and copy number affect methylation rates. (A) Multi-region RRBS was performed on surgically resected non-small cell lung cancers and adjacent normal lung tissue. Sampled regions may differ both in tumor purity and copy number. The observed methylation rate of a differentially methylated CpG is expected to be influenced by purity, copy number and the methylation rate of the normal contaminating cells. (B) Bulk methylation rate histograms for tumor regions 1-3 of patient CRUK0062, for CpGs which are confidently unmethylated in the adjacent normal sample. CpGs are stratified by copy number. A dashed line indicates the expected mode of the methylation rate peak corresponding to clonal differentially methylated CpGs on all copies (***m_t_* = 1**).

### Allele-specific copy number analysis from tumor bisulfite sequencing data

We modeled our approach of purity and copy number inference on existing methods that simultaneously obtain allele-specific copy number and purity estimates from array or sequencing data (Carter et al., 2012; Van Loo et al., 2010; Nik-Zainal et al., 2012). These approaches rely on measures of coverage (LogR) and allelic imbalance (B-allele frequency, BAF) at single nucleotide polymorphisms (SNPs). As CpG islands captured by RRBS are enriched for SNPs (Neininger et al., 2019), we anticipated the data to be highly amenable to allele-specific copy number analysis.

The LogR at SNP loci is readily computed from the normalized read coverage of matched tumor and normal RRBS data. Where patient-matched normal data were not available, the median SNP coverage across all sex-matched normal lung samples in this cohort was used instead. This metric of total copy number suffers from a number of biases in RRBS (**Figure S2**). (i) *MspI* digestion used during library preparation results in a heterogeneous insert size distribution, with fragments ranging from just a few base pairs to hundreds of base pairs in length, depending on the distance between two CCGG recognition sequences (Sun et al., 2015). (ii) Bisulfite conversion alters the GC content of sequences, potentially breaking standard GC-correction for the biases introduced during PCR amplification and Illumina sequencing (Benjamini and Speed, 2012). (iii) Replication timing differs across the genome and between cell types (Ryba et al., 2010). Sequences that replicate early during S-phase tend to have higher coverage than those that replicate later. We correct our LogR estimates for each of these three biases (**Methods**, **Figure S2**).

Due to the bisulfite conversion, compiling reference and alternate allele read counts at SNP loci to obtain BAF values is challenging. Unmethylated cytosines are converted to thymine, yielding four possible bisulfite DNA strands: (complementary to) original top and (complementary to) original bottom (**Figure 2A**). However, most current single-end bisulfite sequencing protocols are directional, yielding only reads from the original top and bottom strands. In addition, as ∼50% of the SNPs captured by RRBS are found at CpG dinucleotides, methylation further complicates BAF calculation.

**Figure 2.**
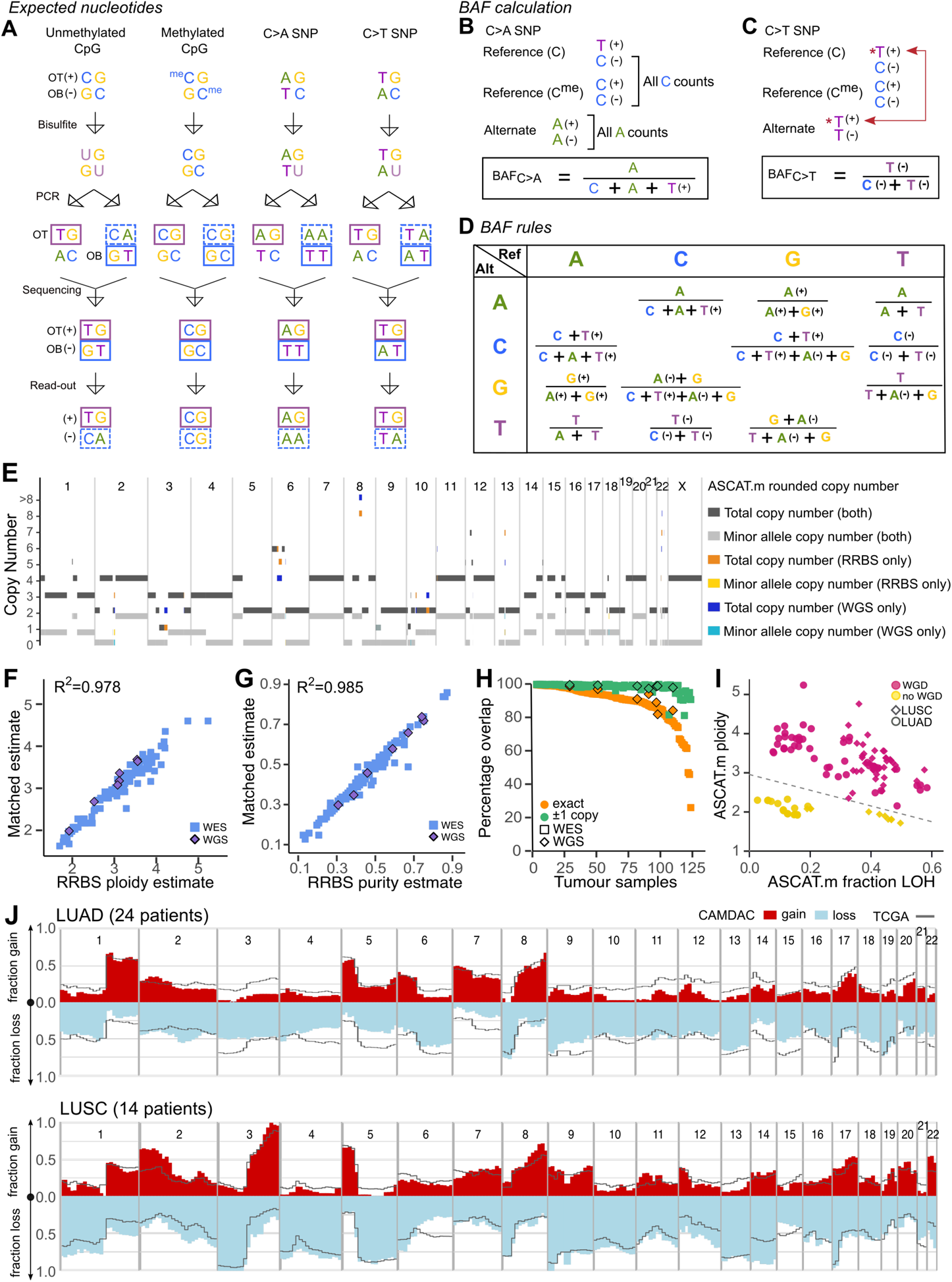
Allele-Specific copy number profiling of RRBS data. (A) Transformation of (un)methylated reference CpG, and alternate ApG and TpG allele dinucleotides during bisulfite sequencing. (B-C) Derivation of BAF rules from strand specific base counts C>A (B) and C>T (C) SNP loci. (D) BAF formulae for all SNP types. (E) Direct comparison of allele-specific copy number estimates derived by ASCAT.m from RRBS and WGS for sample CRUK0069-R1. (F-G) ASCAT.m RRBS-derived ploidy (F) and purity (G) estimates compared with matched whole-exome (WES; blue squares) and, where available, whole-genome sequencing data (WGS; purple diamonds). (H) Percentage overlap between RRBS- and WES- or WGS-derived allele-specific copy number segments. Exact overlaps (orange) and approximate overlaps (±1 copy on a single allele, green) are both displayed. (I) Tumor ploidy and the fraction of the genome with loss of heterozygosity define whole-genome doubling status. (J) Frequency of gains (red) and losses (blue) across the genome of lung adenocarcinoma (LUAD) and squamous cell lung cancer (LUSC). The black line represents gains and losses across 297 LUAD and 444 LUSC samples in TCGA.

Addressing these issues, we propose a set of allele counting and BAF calculation rules for all SNP types, following similar reasoning as Liu et al. (Liu et al., 2012). We illustrate our derivation for C>A and C>T SNPs. Readout bases are reported as original top (+) or complementary to original bottom strand (−) (**Figure 2A-C**). For a C>A SNP at a CpG, all reads reporting A on the + or − strand, i.e. A(+) and A(−), can be uniquely attributed to the alternate allele. Likewise, C(−) derives from the reference allele, together with C(+) and T(+) reads from the methylated and unmethylated cytosine, respectively. As a result, BAF_C>A_ can be calculated as 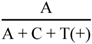 (**Figure 2B**). The situation is more complex for C>T SNPs, since the + strand cannot be used to distinguish between the alternate allele and the bisulfite converted unmethylated reference, as both yield T(+) (**Figure 2C**). However, reads from the − strand do distinguish the (un)methylated reference, C(−), from the alternate allele, T(−). Therefore, it is still possible to quantify BAF_C>T_ as 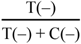. Through similar reasoning, we propose a set of BAF calculation rules for all types of SNPs (**Figure 2D**). The same principles are generalizable to non-CpG cytosine methylation, known to occur at low percentages in the mammalian genome (Schultz et al., 2015). Therefore, our BAF calculation rules are robust to both CpG and non-CpG cytosine methylation. Note that, since only one strand is informative with regard to the allelic imbalance at C>T, T>C, A>G and G>A SNPs, BAF estimates at these sites are based on lower effective sequencing depth.

We validated our approach by comparing genotype calls and BAF estimates at SNP loci on a subset of adjacent normal samples subjected to both RRBS and whole-genome sequencing (WGS, **Methods**). We calculated false positive rates (FPR) and false negative rates (FNR) for SNP calling on RRBS data using the WGS-derived genotypes as ground truth. The average FPR across all SNP types and samples was 0.3% whilst the mean FNR was 25% (**Figure S3A,B**). Polymorphic CCGGs perturbing or creating an *MspI* recognition motif, resulting in allele-specific fragments during RRBS library preparation, and thereby skewing allelic coverage, were the main cause of false negatives (49%) (**Figure S3C,D**). Within the false negative calls, we saw a bias towards SNPs erroneously called homozygous reference (72%) compared to homozygous alternate (28%) (**Figure S3B**), which points towards an alignment bias, likely due to the limited mappability of short *MspI* fragments with alternate alleles.

Leveraging the BAF and LogR values, we infer tumor purity and allele-specific copy number profiles. We call this CAMDAC module ASCAT.m, as our approach relies on ASCAT (Van Loo et al., 2010), but alternative methods such as Battenberg (Nik-Zainal et al., 2012) or ABSOLUTE (Carter et al., 2012) may equally be used. Evaluating ASCAT.m copy number calling by comparing segmented BAF and LogR, and final allele-specific copy number calls to those obtained through WGS of the same tumor samples, confirms the accuracy and robustness of the ASCAT.m estimates (**Figure 2E**).

Comparison of the ASCAT.m RRBS-derived (mean coverage 97x) tumor purity (*ρ*) and ploidy (*ψ*) values with those inferred from WGS (mean coverage 67x) and whole-exome sequencing (WES, mean coverage 464x) performed on the same samples (**Table S3**, Jamal-Hanjani et al., 2017) shows excellent agreement (WGS: *cor*_ρ_ = 0.996, *cor*_ψ_ = 0.991, WES: *cor*_ρ_ = 0.984, *cor*_ψ_ = 0.978) (**Figures 2F-G**). On average, 89.1% of RRBS-derived allele-specific copy number estimates exactly matched those obtained from WES- and WGS data (**Figure 2H**). This percentage increased to 98.4% when allowing a single allele to differ by one copy. Profiling the fraction of the genome with loss of heterozygosity (LOH) and ploidy confirms that whole-genome doubling was a frequent event across the cohort (LUAD: 72%, LUSC: 89%, **Figure 2I**, Dentro et al., 2021; Jamal-Hanjani et al., 2017). In addition, despite low sample sizes, cohort-wide copy-number profiles closely resembled those derived from TCGA LUAD and LUSC cases (**Figure 2J**).

In conclusion, CAMDAC’s ASCAT.m module allows accurate allele-specific copy number profiling and tumor purity estimation from methylation data, obviating the need to perform separate copy number profiling experiments.

### SNP-independent methylation rate estimation

Similar to methylation at CpG SNPs affecting BAF calculations, polymorphisms confound methylation rate estimation (Liu et al., 2012). Polymorphisms at CpGs account for 49.9% of SNPs, 83.3% of which are CpG>TpG polymorphisms, and methylation rates at these positions using standard approaches show markedly different distributions (**Figure 3A-C**). In tumor samples, this effect shows copy number dependence (**Figure 3D-F**).

**Figure 3.**
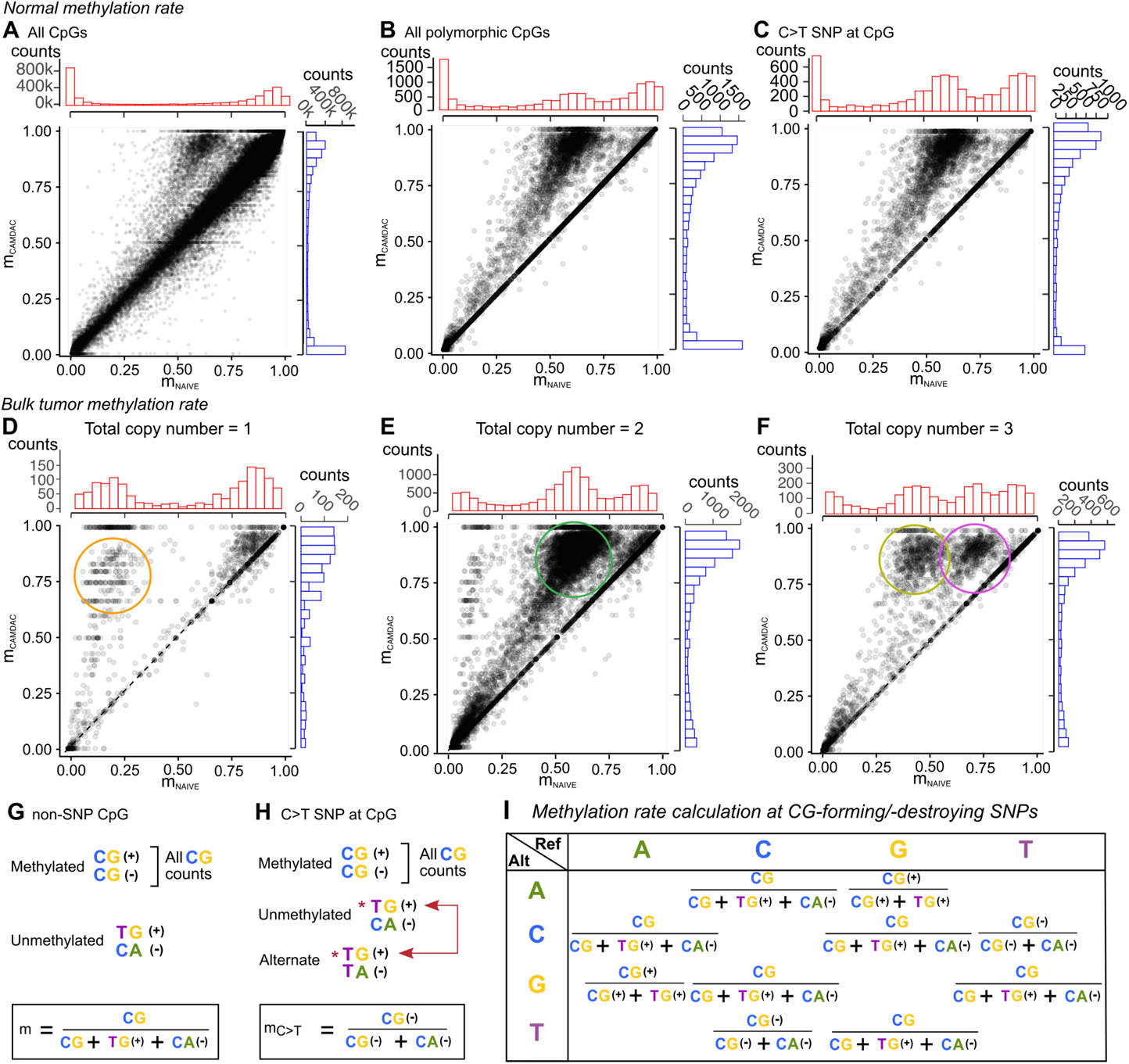
Calculating polymorphism-independent methylation rates with CAMDAC. (A-C) Naïve versus CAMDAC normal methylation rate estimates at (A) all CpGs, (B) polymorphic CpGs and (C) CpG>TpG SNPs, selected from a random sample of 3,000,000 CpGs from this cohort’s 37 normal lung samples. (D-F) Naïve versus CAMDAC bulk tumor methylation rate estimates for CpG>TpG SNPs in segments with total copy number (D) 1, (E) 2 and (F) 3, pooling the data from all 3 sampled regions from patient CRUK0084 of near-equal and high tumor purity (range 0.85-0.87). The data points highlighted by the orange, green and yellow circles indicate heterozygous C>T SNPs with CpG allele copy number 1 and the pink circle CpGs with copy number 2. (G) Derivation of methylation rate estimates at non-polymorphic CpGs. (H) Derivation of the CpG-forming allele-specific methylation rate at a CpG>TpG SNP. (I) Methylation rate formulae for all possible polymorphic CpGs.

To address these biases, we developed an approach to obtain methylation rates independent of SNP genotype (**Figure 3G-I**, **Methods**). A number of considerations need to be made for this. (i) The original top and original bottom strand in directional protocols encode methylation information for the first and second position of a CpG, respectively. As a result, in case of a CpG>TpG SNP, the unmethylated reference and alternate alleles are distinguishable only on the bottom strand. Likewise, at CpG>CpA SNPs, only the top strand may be used. (ii) Computing the methylation rate at the non-polymorphic position in the CpG, only dinucleotides allow to separate the respective contributions from the unmethylated CpG allele and the alternative allele. (iii) To ensure copy number independence of methylation rate estimates, we define them as the average methylation rate per CpG allele. This enables the methylation rate at a heterozygous CpG to vary between 0 and 1, rather than between 0 and 0.5 in a diploid sample, and ensures further independence between methylation rate and copy number estimates. Methylation rates per CpG are thus computed by aggregating strand-specific dinucleotides as follows: 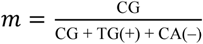, except at C>T and G>A SNPs, where only reads from the bottom strand and top strand, respectively, contribute to the estimates (**Figure 3G-I**).

When accounting for these confounders, the observed biases disappear (**Figures 3A-F**, **S4**), confirming that CAMDAC CpG methylation rate calculation is robust to the presence of polymorphisms.

### Deconvolving tumor methylation profiles

Building upon the allele-specific copy number, tumor purity and methylation rate obtained with CAMDAC, we aim to separate tumor and normal methylation signals. When a tumor clone is methylated at a CpG site that is unmethylated in the admixed normal cells (**Figures S1A-G**), the fraction of methylated reads at that locus observed from bulk bisulfite sequencing increases along with tumor purity and copy number (**Figure 1B**). *Vice versa*, if the tumor clone lost methylation compared to the normal, the observed methylation rate will decrease with increasing purity and copy number (**Figure S1H**). We formally modeled this effect, realizing that the bulk tumor methylation rate (*m_b_*) is the sum of a tumor component (with methylation rate *m*_%_) and a normal component (with methylation rate *m_t_*), weighted by their relative DNA proportions, which can be calculated as the product of purity and copy number (*ρ* and 1 − *ρ*, and *n*_t_ and *n*_n_ for tumor and normal, respectively):

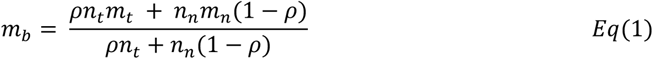

Applying this equation to our non-small cell lung cancer RRBS data, using the adjacent patient-matched normal samples (or a pooled unmatched normal lung reference for sample CRUK0047, **Methods**) as a proxy for the admixed normal cells, enables interpretation of the observed bulk methylation signals. Indeed, the positions of peaks of clonal bi-allelic tumor-normal differential methylation align closely with those predicted under this model, for different values of tumor purity and copy number (**Figures 1B, S1H**), and this relationship holds across all tumor samples (**Figure S5A,B**).

While these data show that our CAMDAC model can account for copy number and tumor purity, not all CpG methylation values appear at the expected peaks. Allele-specific methylation is known to be widespread, not only on the inactive X chromosome in females and at imprinted genes, but also at polymorphic regulatory sequences (Kerkel et al., 2008; Reik and Walter, 2001), and this signal is also apparent in the overall methylation rates (**Figures 1B, S1G-H**). In addition, intra-tumor heterogeneity is expected to contribute intermediate methylation rate signals (Sheffield et al., 2017).

Considering the above, (i) we have formalized the relationship between the bulk and purified tumor methylation rates; (ii) we can directly estimate tumor purity and copy number for each CpG from the bisulfite converted data; and (iii) an adjacent normal sample can be used as a proxy for the methylation rate of the admixed normal cells. We therefore have the necessary information to solve *Eq(1)* for the purified tumor methylation rate:

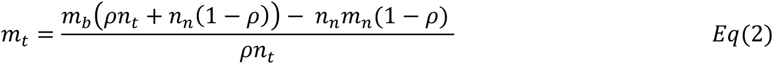

Taken together, bulk tumor methylation rates are confounded by sample purity and local tumor copy number in a way that can be modeled. Formalizing these effects, and inferring purity and copy number from bisulfite sequencing data, CAMDAC corrects bulk methylation rates to yield deconvolved tumor methylation rate estimates.

Post-deconvolution, we observe CpGs that have become methylated on all chromosome copies in all tumor cells to have purified tumor methylation rates approaching the expected maximum *m_t_* = 1. *Vice versa*, tumor-normal hypomethylated loci approach *m_t_* = 0 (**Figure S5C**). Note that the noise on these estimates is proportional to tumor purity and hence statistical power (**Figures S5D-E**).

To evaluate the use of an adjacent normal sample as a proxy for the tumor-admixed normal cell, we used Fluorescent-Activated Cell Sorting (FACS) by DNA content to experimentally separate diploid “normal” cell populations from five bulk tumor samples with clonal whole-genome doubling (average ploidy = 3.53) and performed RRBS on each sorted population (**Methods**). The FACS-purified and adjacent normal methylomes were highly correlated overall (**Figures 4A, S6A**). We next classified CpGs into three groups: (i) confidently unmethylated (UM, HDI^99^ ⊆ [0, 0.2]), (ii) intermediate (IM, HDI^99^ ⊆ [0.2, 0.8]) and (iii) fully methylated (FM, HDI^99^ ⊆ [0.8, 1]) and evaluated correspondence across patient-matched FACS-purified and adjacent normal sample pairs (**Figures 4B, S6B,C**). We observed minimal differential methylation and a gene set enrichment analysis did not reveal any clear gene signatures, suggesting comparatively small differences in cellular composition. These results confirm that in non-small cell lung cancer and at the scale of our biopsies, an adjacent normal sample is an appropriate proxy for the tumor admixed normal.

**Figure 4.**
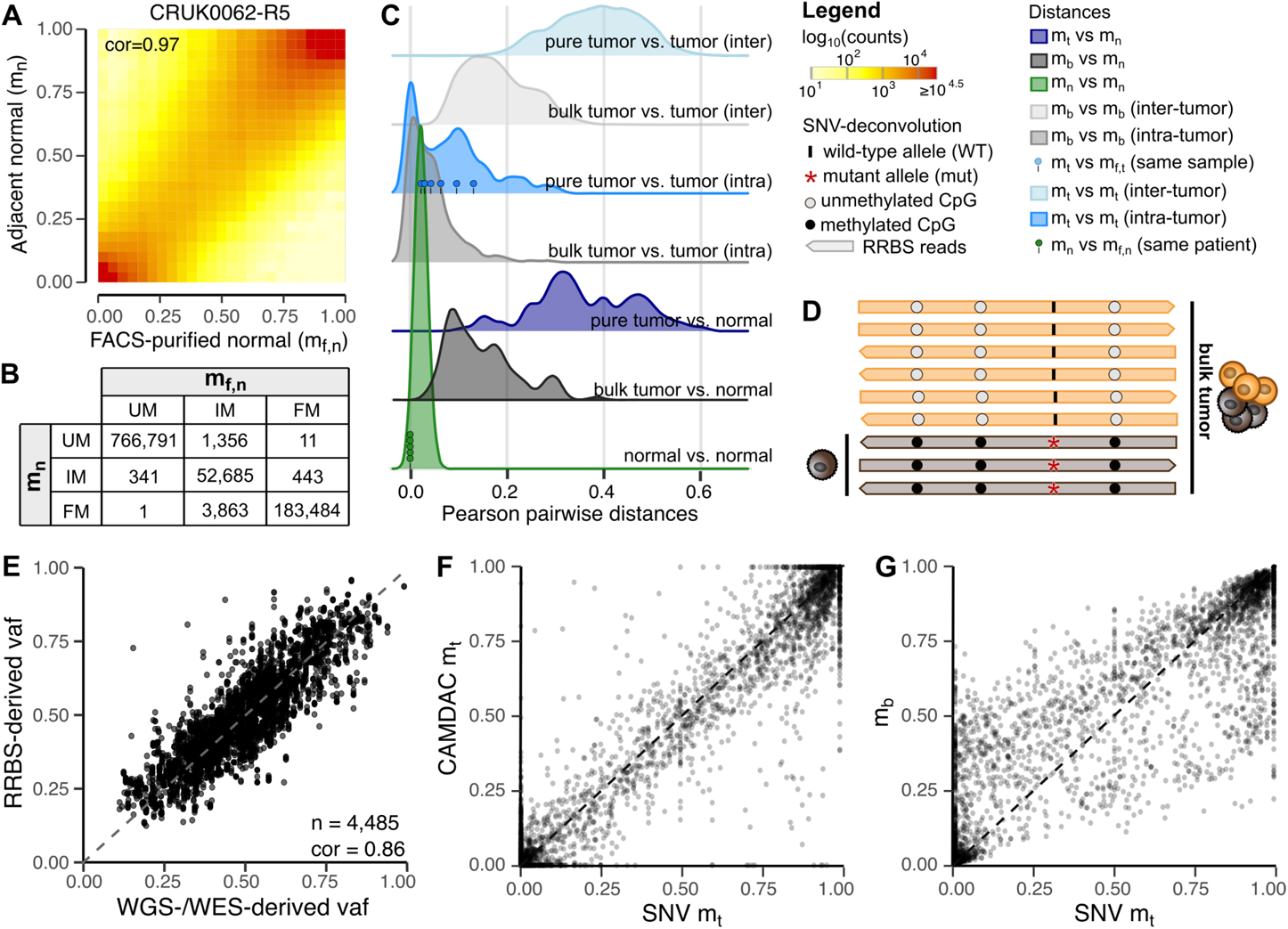
CAMDAC purified methylation profiles. (A) CpG-wise comparison of the adjacent (***m_n_***) and FACS-purified (***m_f,n_***) normal methylation rates with Pearson correlation annotates. (B) CpG classification based on ***m_n_*** and ***m_f,n_***. (C) Correlation between bulk tumor, CAMDAC deconvolved tumor, and adjacent normal methylation profiles. Lollipops represent distances between patient-matched adjacent and FACS-purified normal pairs (green) and between CAMDAC- and FACS-purified tumor methylomes of the same samples (blue). (D) Phasing CpGs to SNVs in regions of LOH enables separation of tumor and normal reads. SNV-purified tumor methylation rates are readily obtained from mutant allele read counts methylated and unmethylated. Remaining reads may be assigned to the wild type allele. (E) Comparison of variant allele frequencies (VAF) of single-nucleotide variants (SNV) derived from RRBS and whole genome/exome sequencing in regions of LOH. (F-G) Validation of CAMDAC methylation rates (F) through phasing of SNVs in regions of LOH and comparison to bulk tumor methylation rates (G).

We evaluated the purified tumor methylation profiles by comparing Pearson distances between pairs of methylation profiles. Compared to bulk signals, CAMDAC deconvolved tumor methylomes show increased distances in inter-patient and tumor-adjacent normal comparisons, while different tumor regions of the same patient remain similar (**Figure 4C**). Distances observed between purified tumor profiles computed with CAMDAC and those obtained experimentally by FACS were low and comparable to those observed between samples taken from the same patient. These observations are in line with admixed normal signals being removed from the bulk tumor and tumor-specific signals being retained in CAMDAC-deconvolved methylation profiles.

Leveraging single nucleotide variant (SNV) calls from additional WGS and previously published WES (Jamal-Hanjani et al., 2017), we validate CAMDAC purified tumor methylation rates by phasing CpG methylation to clonal SNVs present on all chromosome copies in regions with loss of heterozygosity (**Figure 4D**). At these sites, any read reporting the variant allele can directly be assigned to the tumor cells, and methylation rates obtained from this subset of reads should be an unbiased estimate of the purified tumor methylation rate. Overall, RRBS-derived variant allele frequency (VAF) estimates of somatic SNVs (computed analogously to BAF values of germline SNPs, **Figure 2D**) were highly correlated with matched WGS/WES data (Pearson correlation = 0.86, **Figure 4E**). In total, we obtained phased methylation estimates at 4,485 CpG loci across our dataset and observed strong correlation between these SNV-purified *m*_%_ values and CAMDAC estimates (Pearson correlation = 0.97, **Figure 4F**), but not bulk tumor methylation rates (**Figure 4G**). These results confirm that CAMDAC can accurately deconvolute tumor methylation rates from bulk RRBS data.

### Inferring differential methylation from CAMDAC methylation rates

We next set out to formally identify differentially methylated CpGs. For tumor-normal comparisons, we directly compute the probabilities *P*(*m_t_* > *m_n_*) and *P*(*m_t_* < *m_n_*) of hyper- and hypomethylation, respectively. Substituting *m_t_* by *Eq(2)* and modelling the observed bulk and normal methylated read counts in a Bayesian fashion using beta-binomial distributions, the resulting probability density is a scaled difference of two beta posteriors (*m_n_* and *m_b_* ∼ *Beta*(#*reads_meth_*, #*reads_unmeth_*)), which we can compute exactly (**Methods**):

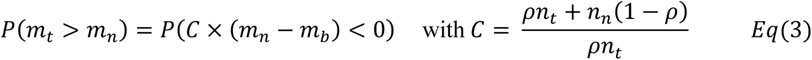

*Equations 2* and *3* reveal how coverage, copy number and tumor purity affect the power to infer tumor–normal differentially methylated positions (DMPs). Directly, the variance of *m_t_* and *m_b_* decreases with increasing normal and tumor coverage, respectively. Indirectly, increasing tumor copy number results in higher local coverage but also, together with increasing purity, shifts *m_b_* away from *m_n_* at DMPs. To identify tumor–tumor DMPs, we similarly compute *P*(*m*_*t*1_ > *m*_%,_) and *P*(*m*_*t*1_ < *m*_%,_) by resampling *m_t_* from the posterior distributions of *m_n_* and *m_b_* in *Eq(2)* (**Methods**).

For differential methylation analysis, it is customary to set a minimal effect size threshold, which we set to Δ*m* > 0.2. In contrast to *m_b_*, setting this threshold using *m_t_* removes dependence on purity and copy number. Simulations based on our observed data (**Figure S7, Methods**) reveal that true positive bi-allelic and mono-allelic DMPs (in balanced regions) indeed show absolute *m_t_* − *m_n_* methylation rate differences near 1 and 0.5 respectively (**Figures 5A and S7D, Methods**). Thresholding on *m_t_* reduces the number of false negative tumor–normal DMP calls in impure tumors and at loci with lower copy number states, while retaining a low false positive rate (**Figures 5B-C and S7E-F**). Similarly, failure to adjust *m_b_* for purity and copy number can readily inundate tumor-tumor differential methylation results with false positive DMPs. Use of CAMDAC *m_t_* values reduces false positives in tumor-tumor comparisons while retaining a similarly low rate of false negatives (**Figures 5D and S7G**).

**Figure 5.**
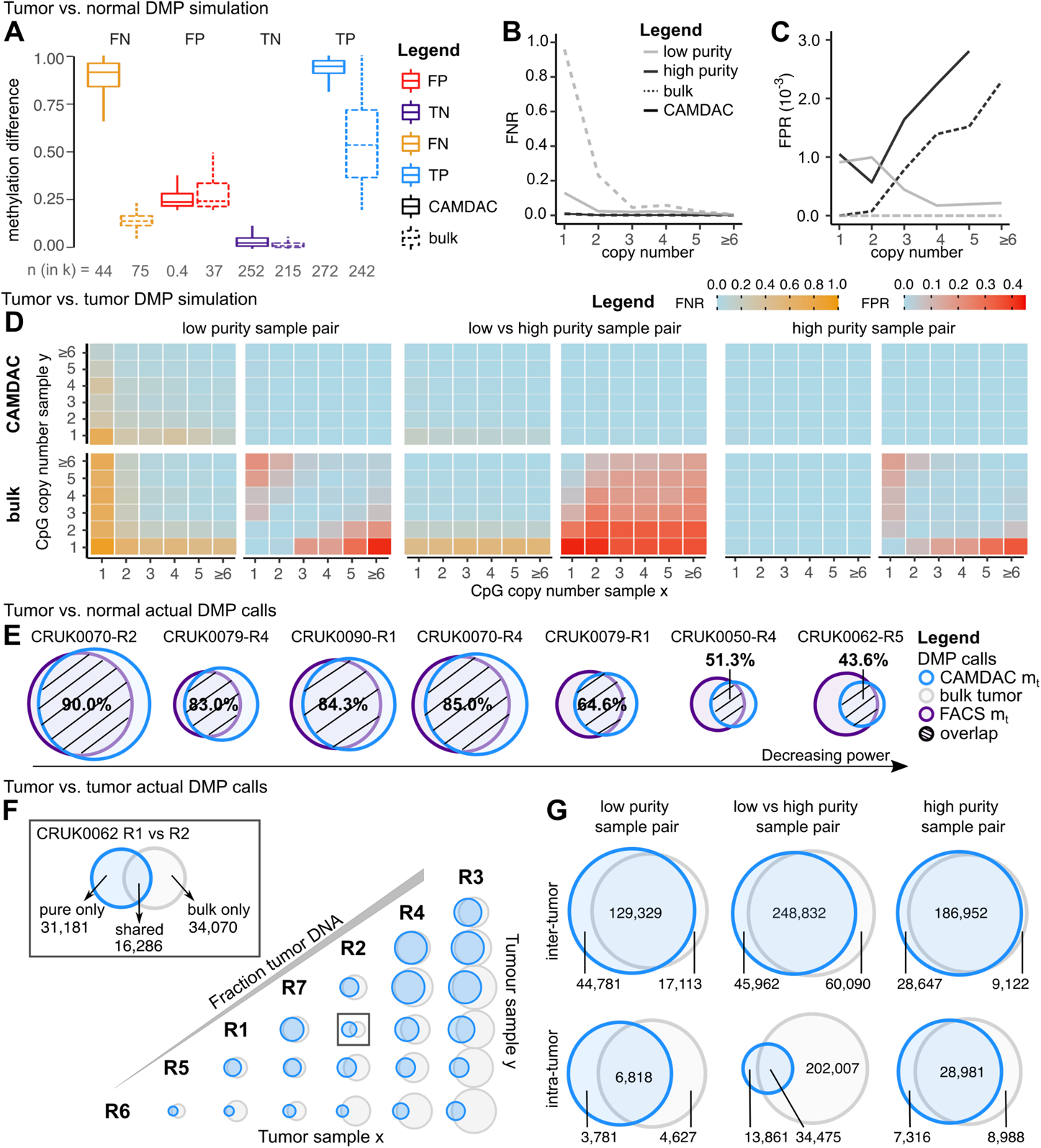
Differential methylation calling using CAMDAC. (A-C) Tumor-normal DMP simulation results. Average methylation difference (A), false negative (B) and false positive rates (C), comparing low versus high simulated tumor purities. (D) Tumor-tumor DMP simulation results. False negative and false positive rates as a function of the copy number state, averaged across simulated sample pairs of low (left panel), low versus high (middle panel) and high (right panel) tumor purities with CAMDAC deconvolved (top row) and bulk (bottom row) tumor methylation rates. (E) Venn diagrams of the overlap between real CAMDAC- and FACS-purified tumor-normal DMPs. The value shown is the percentage of FACS-purified tumor-normal DMPs also called by CAMDAC. Samples are ordered by decreasing power. (F) Comparing observed tumor-tumor differential methylation calls between CRUK0062 regions from bulk (grey) and CAMDAC (blue) approaches. Samples are ordered by fraction tumor DNA content. (G) Venn diagrams showing the overlap of tumor-tumor DMPs between bulk and CAMDAC calls for intra- and inter-tumor DMPs.

To gauge performance on real data, we measured the overlap between tumor-normal DMP calls based on CAMDAC *m_t_* and those obtained from FACS-purified tumor RRBS data of the same tumor regions (*m_f,t_*, **Methods**). While samples were processed from different tissue cuts of the same tumor regions and sequenced to different depths, DMP calls showed 77.3% agreement (**Figure 5E**).

We next compared intra-tumor DMPs called using *m_b_* or *m_t_* for patient CRUK0062, selected for its large number of samples with varying tumor purity (**Figure 5F**). In this setting, 80.9% of CAMDAC *m_t_*-based calls are also identified using *m_b_* and the effect of tumor purity on power can readily be seen as an increase in the number of DMPs identified with sample purity. Using *m_t_*, more DMPs are called when both samples are high purity, *i.e.* statistical power is the highest. In contrast, when using *m_b_*, more DMPs are called when two samples differ more in purity, suggesting the majority are false positives. These findings are in line with our simulation results and suggest that also in real data, controlling methylation rates for tumor purity and copy number greatly reduces the number of false positive DMP calls, while maintaining low false negative rates.

To get a global overview of the performance of CAMDAC *m_t_* and bulk *m_b_* for tumor-tumor differential methylation analysis, we randomly selected CpG loci from samples with a low or high tumor purity both within and between patients and obtained DMP calls (**Methods**). As expected, results showed a greater number of DMPs for inter-patient comparisons than between samples of shared clonal origins (**Figure 5G**). Furthermore, DMP calls unique to the bulk tumor were frequent between samples of differing purities taken from the same patient, in line with the high false-positive rate of DMP calling without deconvolution.

When investigating disease-related DNA methylation changes, researchers commonly look for differentially methylated regions (DMRs) as opposed to individual CpGs (Robinson et al., 2014). Building on DMP calls, CAMDAC can identify DMRs by binning CpGs into neighborhoods and identifying DMP hotspots within these clusters (**Methods**).

Taken together, analyses of both simulated and observed data show that CAMDAC enables accurate calling of both tumor-normal and tumor-tumor differential methylation from RRBS data.

### Quantifying allele-specific methylation

While most CpGs are either fully methylated or fully unmethylated in the tumor, others show intermediate *m_t_* values (**Figures 1B, 4G, S1G**). We hypothesize that, in addition to intra-tumor epigenetic heterogeneity, part of this is due to allele-specific differential methylation, and we set out to quantify this phenomenon. Note that, by construction, allele-specific methylation at heterozygous CpGs does not contribute to this intermediate *m_t_* signal (**Figure 3H-I**).

To assess CAMDAC sensitivity for intermediate methylation signals, we first evaluated methylation rates at the imprinted *IGF2/H19* locus (Frevel et al., 1999). As expected, the normal methylation rate around the imprinting control region is confidently around *m_t_*= 0.5, whilst the CpG island shore is fully methylated (e.g. CRUK0073, **Figure 6A**). Since imprinting results in allele-specific methylation, we aimed to validate this across known imprinted loci (Geneimprint database, **Methods**) by phasing CpGs to nearby germline heterozygous SNPs. In total, we detected allele-specific methylation at 100/106 (94%) loci in normal samples with phased methylation rates and leveraged the CAMDAC equations to deconvolve allele-specific tumor methylation rates (**Methods**).

**Figure 6.**
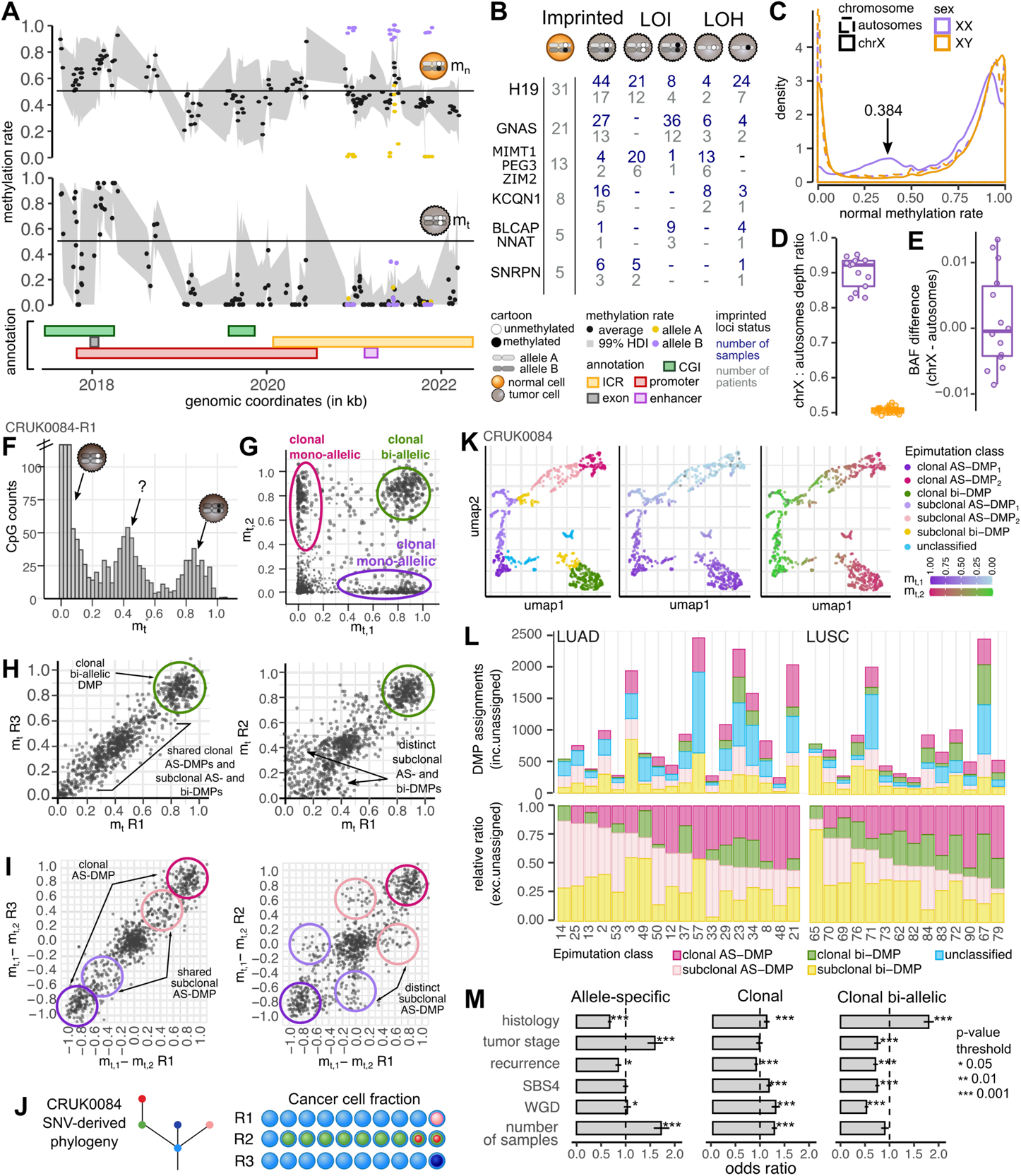
Quantifying allele-specific methylation. (A) CRUK0073 normal (CRUK0073-N, top) and CAMDAC purified tumor (CRUK0073-R2, middle) CpG methylation rate along the H19 locus with methylation rate point estimates shown as black dots, HDI_99_ displayed as a grey ribbon and genomic annotations showing the transcript and promoter region as well as neighboring CpG islands, enhancers and the imprinting control region (ICR, bottom panel). Phased methylation rates to both parental alleles are shown (purple and yellow). (B) Overview of somatic changes at known imprinted loci with phased methylation information across tumor samples. LOI: loss of imprinting; LOH: loss of heterozygosity. (C) Allele-specific methylation signals on chromosome X in adjacent normal lung samples across female and male patients. (D, E) Comparison of (D) sequencing depth and (E) mean BAF at heterozygous SNPs on chromosome X to that on autosomes in males and females. (F) Deconvolved pure tumor methylation rates from CRUK0084-R1, including only phaseable CpGs which are confidently unmethylated in the matched adjacent normal sample and located on segments with allele-specific copy number 1+1. (G) Scatter plot of the allele-specific pure tumor methylation rates of the CpGs loci in (F). (H-I) Comparison of (H) pure tumor methylation rate differences and (I) average pure tumor methylation rates between alleles across samples pairs for the same patient reveals subclonal relationships. (J) SNV-based phylogenetic clone tree for CRUK0084 taken from Jamal-Hanjani et *al*. (Jamal-Hanjani et al., 2017). (K) UMAP of allele-specific pure tumor methylation rates including CpGs that are hypermethylated in at least 1 sample. Hierarchical clustering and analysis of cluster methylation rates enables epimutation assignments, taking copy number into account. (L) Total number of unique epimutations per class (top panel) and their ratio (bottom panel) per patient, colored by clonality and allele-specificity. (M) Allele-specific (left) and clonal (middle) and clonal bi-allelic only (right) DMP odds ratio given clinical features including histological subtypes (LUAD and LUSC), stage (1a and 1b against later stages) and genomic annotations, including whole genome doubling status and smoking signature SBS4 exposure (high > 0.35, low otherwise) and number of samples sequenced (high > 3, low otherwise).

In line with previous reports (Cui et al., 2002; Ogawa et al., 1993), we observed frequent loss of imprinting in tumors and report cases with or without loss of heterozygosity, accounting for 74/276 (26.8%) and 102/276 (37.0%) loci, respectively (**Figure 6A,B**). Loss of imprinting without loss of heterozygosity showed a preference for both parental alleles losing (*IGF2/H19*, *MIMT1/ZIM2/PEG3*, *SRNPN*) or gaining methylation (*GNAS/GNASAS*, *BLCAP/NNAT*, **Figure 6B**). Overall, somatic alterations at imprinted loci were frequently ubiquitous among samples from the same patient (39/60, 65%, **Figure S8A**). Interestingly, we report 7 cases with distinct subclonal alterations affecting the same locus. We map these epimutations to phylogenies of the same samples published in Jamal-Hanjani *et al*. (Jamal-Hanjani et al., 2017), and find evidence of 4 parallel and 2 sequential evolutionary events (**Figure S8B**). We therefore posit that deregulation of imprinted loci may play a role in shaping a subset of non-small cell lung cancers.

Next, we evaluated allele-specific methylation signals in adjacent normal samples from females. Dosage compensation in females by X-chromosome inactivation involves extensive DNA methylation of promoter-associated CpGs on the inactive copy (Lyon, 1962). Unexpectedly, intermediate methylation values on X were consistently below 0.5 (median *m_t_* = 0.384) for the 13 normal female samples in this cohort (**Figure 6C**). We observed decreased sequencing coverage on X compared to the rest of the genome in females (**Figure 6D**) while BAF of heterozygous SNP loci did not show any deviation from 0.5 (**Figure 6E**). As this effect was absent from WES data (data not shown), this suggests that the RRBS library preparation may be biased against the methylated Barr body. Due to the mosaic nature of X chromosome inactivation in females, at the scale of our adjacent normal bulk lung samples, both parental X chromosomes are inactivated in an approximately equal number of cells, and no enrichment of allele-specific methylation on chromosome X was observed compared with autosomes (t-test, p = 6.36 × 10^−2^).

Leveraging allele-specific pure tumor methylation rates at CpG loci that were confidently unmethylated in the matched normal (HDI_99_ [0, 0.2], **Methods**), we demonstrate that allele-specific CpG hypermethylation contributes to intermediate *m_t_* values and confirm the presence of bi-allelic alterations (**Figure 6F,G**). In addition, direct comparison of average and allele-specific methylation rates at the same CpGs between sample pairs allows visualization of (sub)clonal allele-specific methylation clusters (**Figures 6H**). These clusters recapitulate the previously reported SNV-derived phylogeny of the same tumor, but also enable detection of further subclones (**Figure 6I,J**). Through uniform manifold approximation and projection (UMAP) analysis of allele-specific and average pure tumor methylation rates across samples from the same patient, followed by clustering, we classified epimutations as (sub)clonal bi-allelic and mono-allelic (**Methods**, **Figure 6K**).

Interestingly, while varying widely for individual tumors, we observed that the extent of allele-specific *vs.* bi-allelic and subclonal *vs.* clonal methylation changes differed between LUSC and LUAD and stage I *vs.* stage II/III tumors (**Figure 6L,M**). This suggests distinct roles and mechanisms underlying DNA methylation heterogeneity and evolution in these tumor types. The fraction of clonal bi-allelic DMPs was the strongest indicator of relapse and could present a unique opportunity for early patient stratification.

### Interplay of somatic mutations and methylation changes

To gain insight into the relationship between somatic mutations and epimutations, we next performed DMR calling on the SNV-purified tumor methylation rate estimates. In total, 4,406 methylation bins comprised of 3 or more CpGs could be phased to SNVs, of which 610 were differentially methylated (**Figure 7A**, **Table S4**). These SNV-phased DMRs were more frequently hyper-than hypomethylated, potentially resulting from the disruption of transcription factor binding sites and thus decreased enhancer binding affinity (Morova et al., 2020; Zhao et al., 2017).

**Figure 7.**
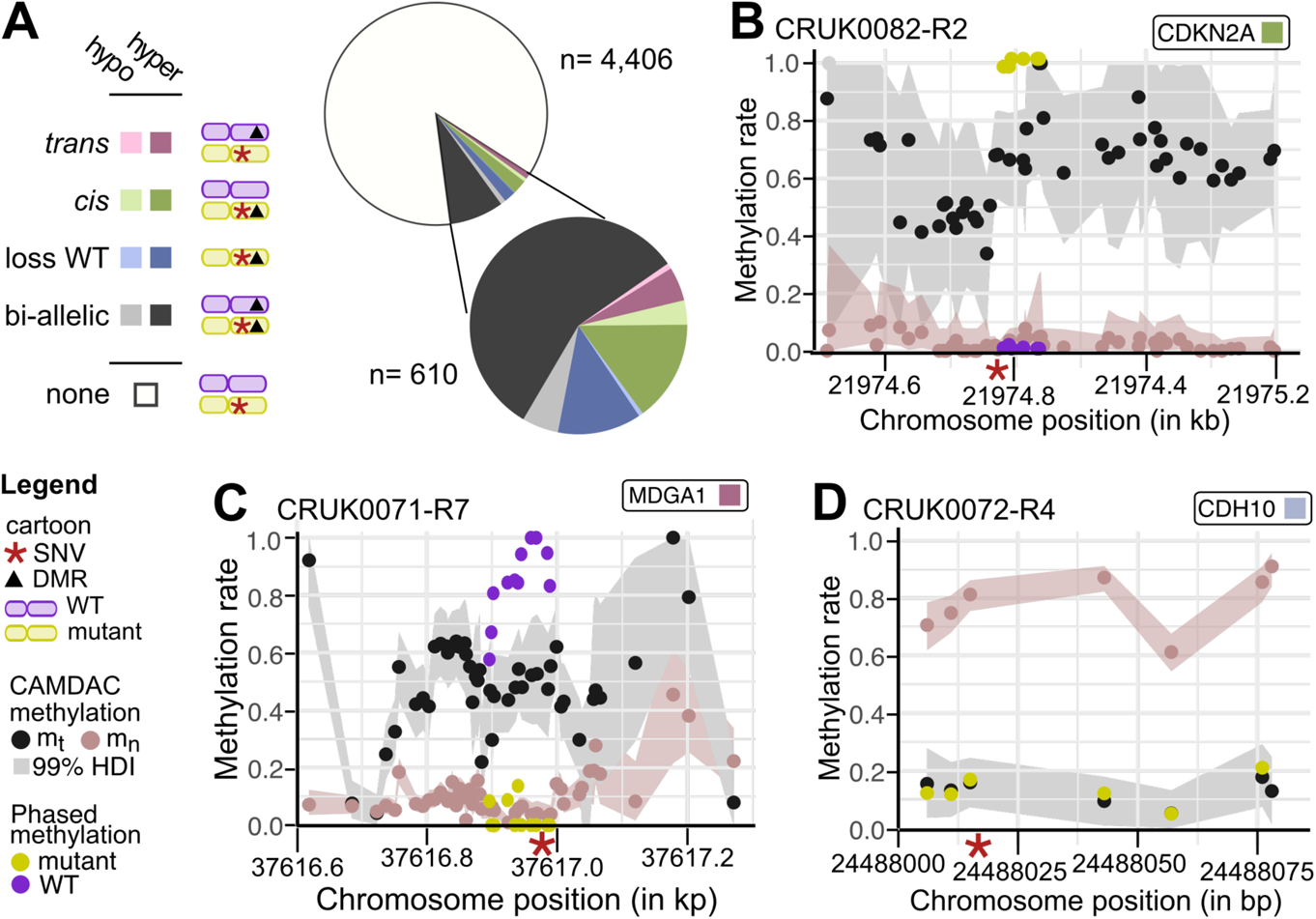
The interplay of somatic mutations and methylation changes. (A) Overview of phasing results of 4,406 methylation bins to somatic SNVs. DMRs are annotated as either in-*trans* or in-*cis* to the SNV, on both alleles, or showing loss of the wild type (WT) allele. Each DMR category is further sub-divided into hyper- and hypomethylated subgroups. (B-D) Examples of SNV-phased, CAMDAC deconvolved tumor and matched normal methylation rates across genes harboring SNVs and DMRs in-*cis* (B), in-*trans* (C), and with loss of the WT allele (D).

Supporting the functional consequences of mutations and/or increased mutability of methylated cytosines, 114 *vs.* 36 hypermethylated SNV-phased DMRs without loss of the wild type allele were specific to the mutant (in-*cis*) and the wild-type (in-*trans*) allele, respectively (p-val = 1.22 × 10^−10^, two-sided Binomial test). For example, patient CRUK0082 harbors a clonal *CDKN2A* promoter mutation, where, in regions with phased methylation information, only the mutant allele is hypermethylated (**Figure 7B**).

In-*trans* hypermethylation events may occasionally constitute inactivation of genes due to deleterious mutation of one allele and hypermethylation of the other. In one potential example, enhancer GH06F037648, which is intragenic to *MDGA1*, was found to be clonally hypermethylated on one copy of the wild type allele in all regions from patient CRUK0071 (**Figure 7C**).

In regions where the wild type allele is lost, it is not possible to determine whether in-*cis* DMRs were originally restricted to the mutant allele and thus potentially linked to the genetic alterations, or on all copies, suggesting the two events were independent. As an example of this, we observe clonal loss of the wild type allele, combined with a missense mutation and hypomethylation of surrounding CpGs at the *CDH10* locus in patient CRUK0072 (**Figure 7D**).

Taken together, CAMDAC deconvolution and phasing enables deeper understanding of the interplay between aberrant DNA methylation and genetic mutations. In our non-small cell lung cancer cohort, we observed frequent phasing of the hypermethylated allele to the mutant SNV allele, potentially through ablation of adjacent transcription factor binding sites.

### Relationships between purified methylation profiles

To gain insight into similarities and differences between samples, we clustered genome-wide methylation profiles. We focused on tumor-normal promoter DMRs, selected for being enriched in gene regulatory driven differential methylation, and performed UMAP of deconvolved tumor and normal methylation rate profiles (**Figures 8A,B**). We identified four main clusters: two clusters of cancer samples and two clusters of adjacent normal samples. Normal lung epithelium samples clustered by sex (**Figure 8A**), indicating that differences are dominated by female-to-male differential methylation. These sex-based differences were less dominant in the tumor samples, likely in part because many female samples showed complete (34%) or at least partial (52%) LOH of chromosome X. In contrast, non-small cell lung cancer samples mostly separated by histology (**Figure 8B**), pointing towards large methylation differences between adenocarcinoma and squamous cell carcinoma, possibly reflecting a different cell of origin (Sutherland and Berns, 2010). Different tumor samples from the same patient consistently clustered together (**Figure 8C,D**).

**Figure 8.**
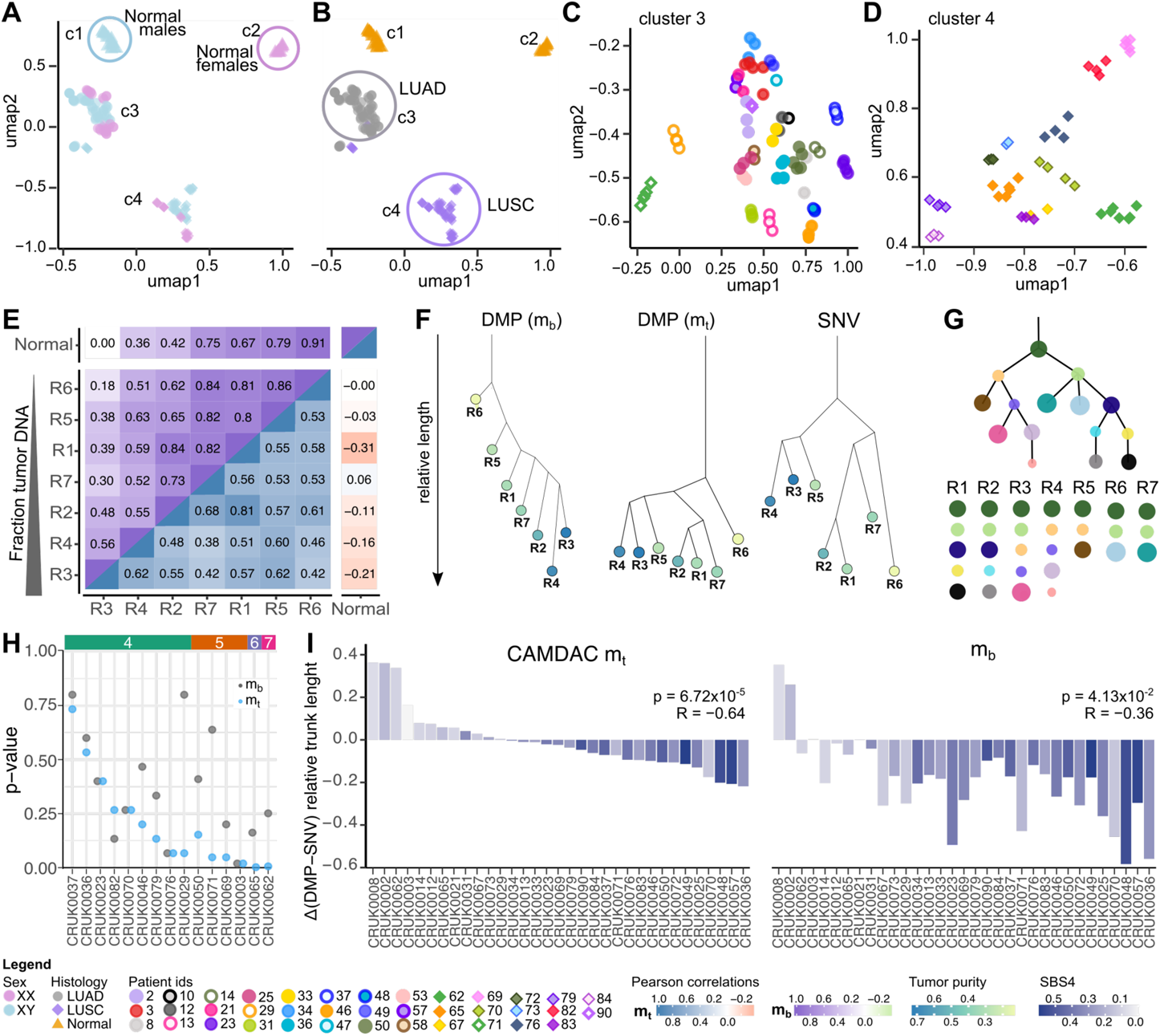
CAMDAC deconvolved tumor methylation rates inform clustering and intra-tumor heterogeneity analyses. (A-B) UMAP of the mean purified tumor methylation rates at CpGs in tumor-normal promoter DMRs, (A-B) annotated with sex (A) and sample histology (B). (C-D) Zoom in of UMAP clusters 3 (C) and 4 (D), annotated with patient IDs. (E) Correlations between bulk (upper triangle) and purified methylation rates (lower triangle) and matched normal (upmost and right-most columns) using all tumor-normal DMPs in one or more of the seven tumor samples from CRUK0062. Samples are ordered by tumor purity. (F) Neighbor-joining phylogenetic sample tree for CRUK0062 based on binarized bulk tumor and CAMDAC deconvolved tumor methylation estimates at tumor-normal DMPs or using binarized CCF at SNVs. (G) SNV-based phylogenetic clone tree for CRUK0062 from Jamal-Hanjani et *al*. (Jamal-Hanjani et al., 2017). (H) Empirical p-value obtained when fitting (hyper)methylation data to the matched SNV tree for each patient. Patients are grouped by the number of sampled tumor regions with RRBS data (excluding the adjacent normal). (I) Difference in the trunk to longest branch length ratio between DMP and matched SNV sample trees. Left: CAMDAC pure tumor DMP *vs.* SNV sample trees; Right: bulk tumor DMP *vs.* SNV sample trees.

### Purified methylation profiles reveal phylogenetic relationships

We next focused more closely on the relationship between methylation profiles of different samples from the same patient. For each patient, we select CpG sites that are differentially methylated compared to the normal in at least one sample. After deconvolution, tumor signals show reduced correlation with adjacent normal methylation rates, and tumor purity is no longer the main driver of inter-sample correlations (**Figure 8E**).

Instead, CAMDAC pure tumor methylation profiles may encode evolutionary relationships between subclones. Indeed, sample phylogenetic trees constructed from methylation rates at (hypermethylated) tumor-normal DMPs recapitulate clonal phylogenetic relationships inferred from matched WES (Jamal-Hanjani et al., 2017, **Methods**, **Figures 8F,G and S9**). In contrast, bulk methylation profiles were unable to reproduce these relationships, and generally clustered by tumor purity. Overall, methylation trees based on CAMDAC *m_t_* outperformed *m_b_* in reproducing phylogenetic relationships (combined empirical p-values = 1.61 × 10^−5^ and 0.109, respectively, Fisher’s method, **Figures 8H and S10**, **Methods**).

Epigenetic alterations typically outnumber somatic mutations, and may accumulate early in tumor evolution (Baylin and Jones, 2016), creating a permissive state for tumor initiation. As such, we expected longer trunk lengths (relative to subclonal branches) in DMP-than SNV-based patient trees. We calculated the relative trunk length difference between DMP and SNV tree pairs and saw longer trunks in CAMDAC *m_t_*-derived DMP trees than in SNV trees, but only in patients with low SBS4 smoking signature exposure (Pearson correlation = −0.64, p-value = 6.72 × 10^−5^, **Methods**, **Figure 8I**). This trend was lost in the bulk tumor data, where virtually all DMP trees had shorter trunks compared to SNV trees due to signal from normal contaminating cells.

We therefore conclude that CAMDAC provides unique opportunities to study intra-tumor heterogeneity in solid tumors, unconfounded by signals from admixed normal cells.

## DISCUSSION

Bulk tumor methylation signals are confounded by copy number aberrations and normal cell admixture. CAMDAC determines tumor purity and allele-specific copy number from bulk tumor-normal matched RRBS data and uses this to reconstruct the true tumor (allele-specific) methylation rate from the bulk tumor and matched normal rates. Use of these purified methylomes increases the accuracy of tumor-normal and tumor-tumor differential methylation calling and can reveal the phylogenetic relationships between subclones. In addition, they provide insight into tumor biology and the taxonomy of cancer.

While a few studies have considered either the effect of sample purity (Hua et al., 2020; Zheng et al., 2017) or copy number (Martin-Trujillo et al., 2017) on bulk tumor methylation rates, CAMDAC is – to our knowledge – the first approach to estimate and leverage both confounders to reconstruct the tumor methylation signal. We demonstrate that tumor-normal differential methylation produces predictable intermediate methylation values given local copy number states. This explains, at least in part, why bulk cancer methylomes exhibit an increased proportion of intermediately methylated regions compared to normal tissues (Lister et al., 2009).

After purification, most differentially methylated CpGs are either fully methylated or unmethylated. Nevertheless, a significant subset remains intermediately methylated. Allele-specific CAMDAC-purified tumor methylation rates allow us to decipher this signal and classify epimutations as (sub)clonal bi- or mono-allelic. From these data, we suggest two potential classes of DNA methylation changes: (i) gene regulatory differential methylation and (ii) allele- (and copy-) specific stochastic methylation changes. Regulatory changes typically affect all copies indiscriminately, with possible exceptions at heterozygous loci, while stochastic methylation changes arise due to errors in the DNA methylation maintenance machinery, aberrant TET activity, or spontaneous deamination of methylated cytosines (Coulondre et al., 1978; Goyal, 2006; Kohli and Zhang, 2013). Assuming faithful replication thereafter, these changes become discernable after clonal expansion, generating intermediate methylation signals.

CAMDAC methylation profiles recapitulate the phylogenetic relationships between subclones and unlock a wealth of information on intra-tumor heterogeneity. CAMDAC also recovers the clonal DMP signals that are lost in bulk methylation data due to normal cell contamination. Additionally, CAMDAC DMP trees of both LUSC and LUAD cases had longer relative trunk lengths than SNV trees in the absence of highly mutagenic processes such as smoking, which suggests that DNA methylation may have a role in early tumor evolution. We envisage further development will enable reconstruction of subclones within single samples and accurate clone trees (as opposed to sample trees, Alves et al., 2017) from bisulfite sequencing data, akin to current methods leveraging single-nucleotide variants (Dentro et al., 2017; Nik-Zainal et al., 2012; Tarabichi et al., 2021). As such, application of CAMDAC will help elucidate the role of epimutations in the evolutionary histories of different cancer types.

As in previous studies (Gaiti et al., 2019; Hua et al., 2020), we infer differential methylation compared to adjacent normal tissue and use this as a proxy for the admixed normal cells in the tumor sample. Where matched normal tissue samples are not available, a tissue-matched normal reference may be constructed for use with CAMDAC. Outside our cohort, the DNA methylation data flowing from large-scale initiatives such as the International Human Epigenome Consortium (IHEC, Stunnenberg et al., 2016) and the ENCODE (Feingold et al., 2004) and BLUEPRINT (Fernández et al., 2016) projects, enable construction of reference profiles for a wide range of tissues.

In low purity samples and in regions with lower tumor copy number, fewer reads report on the methylation state of the tumor. CAMDAC incorporates this greater uncertainty in its estimates for the tumor methylation rates in such cases, greatly reducing the number of false positive hits when inferring differential methylation. Bulk tumor methylomes of low purity samples are dominated by signals from the admixed normal cells, as evidenced by the high false negative rate at simulated tumor-normal epimutations based on bulk profiles. In tumor-tumor differential methylation analyses based on bulk tumor methylomes, more DMPs are identified when two samples differ more in purity, suggesting a high false positive rate. In contrast, when using CAMDAC pure tumor methylation rate increases, the number of DMPs increases with statistical power.

DNA methylation heterogeneity has been linked to clonal progression in several tumor types (Hao et al., 2016; Mazor et al., 2015). Various metrics have been conceived to quantify intra-tumor methylation heterogeneity from bulk data and some have shown prognostic value in leukemia and Ewing sarcoma (Landau et al., 2014; Li et al., 2016; Sheffield et al., 2017), but not in glioblastoma (Klughammer et al., 2018). While heterogeneity estimates from bulk tumor methylomes are likely accurate in tumor types for which high purity samples are readily obtained, we hypothesize these metrics are similarly confounded by normal cell admixture and copy number aberrations in most other cases. Applying CAMDAC principles to adjust these metrics accordingly may yield powerful prognosticators and shed further light on the role of DNA methylation heterogeneity in solid tumors.

In summary, CAMDAC enables us to mine a wealth of information contained in bulk tumor bisulfite sequencing data, combining purity and copy number profiling with accurate differential methylation analysis based on the purified tumor methylation rate estimates. CAMDAC uniquely unlocks insights into intra-tumor heterogeneity and the biology and taxonomy of cancer. We expect CAMDAC will further our understanding of the interplay between genetic and epigenetic mutations and their roles in tumor evolution.

## ACKNOWLEDGMENTS

This work was supported by the Francis Crick Institute, which receives its core funding from Cancer Research UK (FC001202), the UK Medical Research Council (FC001202), and the Wellcome Trust (FC001202). For the purpose of open access, the authors have applied a CC BY public copyright licence to any author accepted manuscript version arising from this submission. We acknowledge technical support from the CRUK–UCL Centre-funded Genomics and Genome Engineering Core Facility of the UCL Cancer Institute and grant support from the NIHR BRC (BRC275/CN/SB/101330) and the Wellcome Trust (218274/Z/19/Z). J.D. is a postdoctoral fellow supported by the European Union’s Horizon 2020 research and innovation program (Marie Skłodowska-Curie Grant Agreement No. 703594-DECODE) and the Research Foundation – Flanders (FWO; 12J6921N). M.T. was supported by the People Programme, Marie Skłodowska-Curie Actions (FP7/2007-2013/WHRI-ACADEMY-608765) and the Danish Council for Strategic Research (1309-00006B). A.F. received funding from Prostate Cancer UK (matr15-009), Biotechnology and Biological Sciences Research Council (BB/R009295/1) and the Medical Research Council (MR/M025411/1). C.S. is Royal Society Napier Research Professor (RP150154). His work is supported by the Francis Crick Institute, which receives its core funding from Cancer Research UK (FC001169), the UK Medical Research Council (FC001169), and the Wellcome Trust (FC001169). C.S. is funded by Cancer Research UK (TRACERx, C11496/A17786; Cancer Research UK Lung Cancer Centre of Excellence C11496/A30025), the Rosetrees Trust, Butterfield and Stoneygate Trusts, NovoNordisk Foundation (ID16584), a Royal Society Research Professorship Enhancement Award (RP/EA/180007), the National Institute for Health Research (NIHR) Biomedical Research Centre at University College London Hospitals, the Breast Cancer Research Foundation (BCRF 20-157), the CRUK-UCL Centre, Experimental Cancer Medicine Centre. His research is supported by a Stand Up To Cancer-LUNGevity-American Lung Association Lung Cancer Interception Dream Team Translational Research Grant (SU2C-AACR-DT23-17). C.S. also receives funding from the European Research Council (FP7-THESEUS-617844, FP7-PloidyNet 607722, PROTEUS 835297 and Chromavision 665233). M.J.-H. is a CRUK Career Establishment Awardee and has received funding from CRUK, IASLC International Lung Cancer Foundation, Lung Cancer Research Foundation, Rosetrees Trust, UKI NETs, NIHR UCLH Biomedical Research Centre. P.V.L. is a Winton Group Leader in recognition of the Winton Charitable Foundation’s support towards the establishment of The Francis Crick Institute. P.V.L. is a CPRIT Scholar in Cancer Research and acknowledges CPRIT grant support (RR210006).

## AUTHOR CONTRIBUTIONS

E.L.C. developed ASCAT.m and CAMDAC, under the supervision of J.D. and P.V.L, with important inputs from S.B.; J.D. conceptualized CAMDAC. C.S. and S.B. conceptualized the TRACERx methylation study and acquired financial support, with input from G.A.W., A.F., and M.T.; M.T., G.A.W., H.V., A.F. and P.D. set up and processed the bulk RRBS. A.V. and C.C.M. performed FACS analysis followed by RRBS, with input from N.K. and assistance from the cell sorting and sequencing facilities at the Francis Crick Institute. E.L.C. performed analyses with contributions from N.E.M. and C.C., and input from T.B.; E.L.C. visualized data and designed figures, with additions from N.E.M. and C.C.; T.B.K.M. and M.D. performed WGS analyses under the supervision of N.M. and C.S.; S.V., M.J.H. and C.S. were critical in the funding and acquisition of clinical samples.; J.D. and P.V.L. supervised the study. E.L.C., J.D. and P.V.L. wrote the manuscript, with significant contributions from N.E.M. and C.C., and further input from S.B., M.T. and C.S. All authors read and approved the final manuscript.

## SUPPLEMENTARY FIGURES

**Figure S1.**
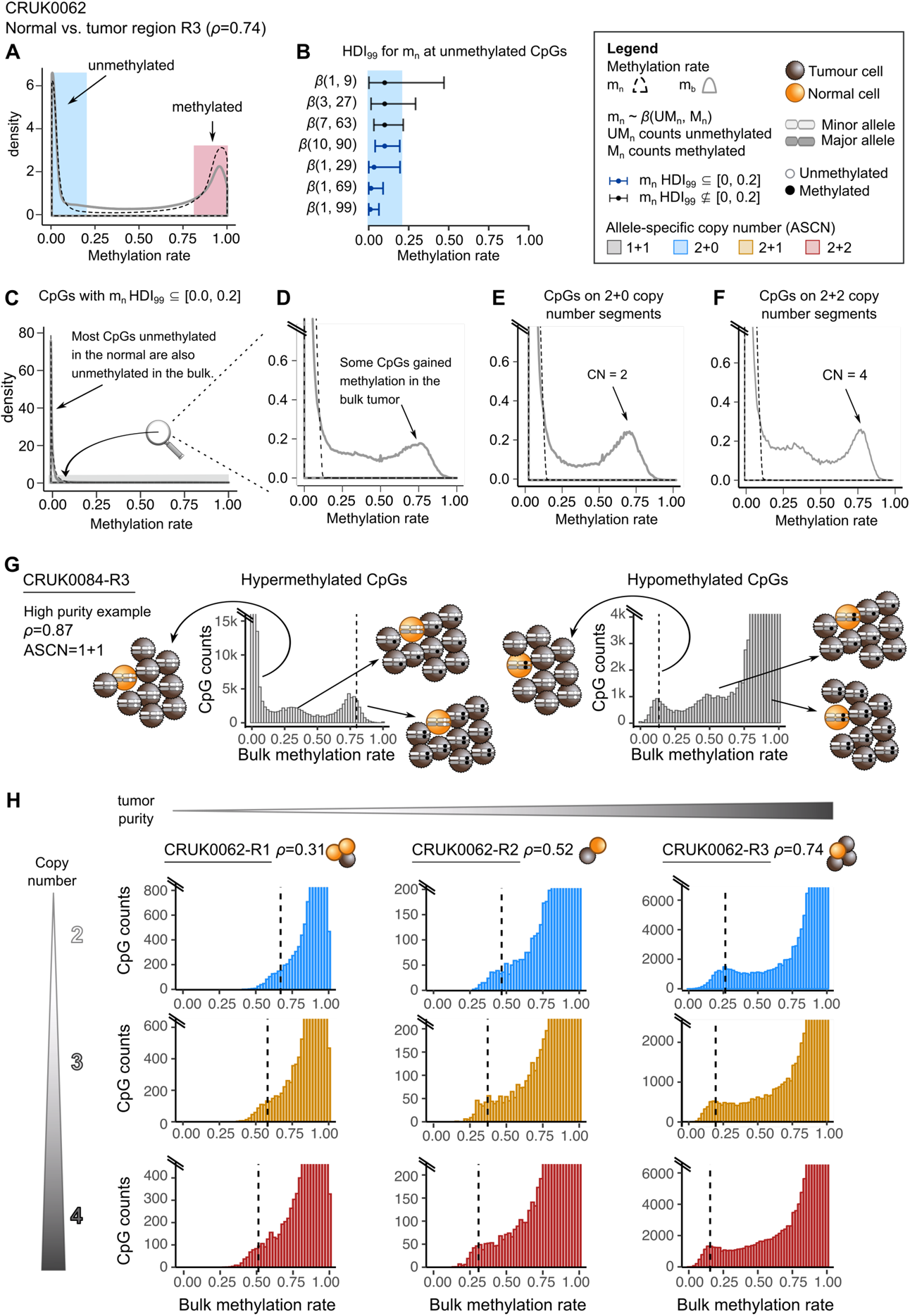
Tumor purity and copy number affect methylation rates. (A) Normal and bulk tumor methylation rate density for sample CRUK0062-R3 and adjacent patient-matched normal showing most CpGs are either fully methylated or unmethylated. An increased number of CpGs with intermediate methylation levels is found in the bulk tumor. (B) Simulated 99% Highest Density Interval (HDI_99_) at example unmethylated CpGs with variable number of methylated and unmethylated read counts and total CpG coverage. (C-F) Normal and bulk tumor methylation rate (*m_t_* and *m_b_*, respectively) density plot for CpGs labelled as confidently unmethylated in the normal (HDI_99_ ⊆ [0,0.2]). The majority of these loci are also unmethylated in the bulk tumor. (D-F) Zoom-in, highlighting CpGs with gained methylation in the tumor cells across all copy numbers (D), and in regions of copy number states 2+0 (E) and 2+2 (F). (G) Bulk tumor methylation rate histograms of CRUK0084-R3, having an estimated tumor purity (*ρ*) of 87%, for CpGs in diploid regions confidently unmethylated (left) and methylated (right) in the patient-matched adjacent normal. Hyper- and hypomethylated CpGs can be found in peaks that vary with the mutation copy number. (H) Bulk methylation rate histograms of CRUK0062 tumor regions 1-3, for CpGs which are confidently methylated in the adjacent normal sample, stratified by allele-specific copy number state. A dashed line indicates the expected mode of the methylation rate peak corresponding to clonal differentially methylated CpGs on all copies.

**Figure S2.**
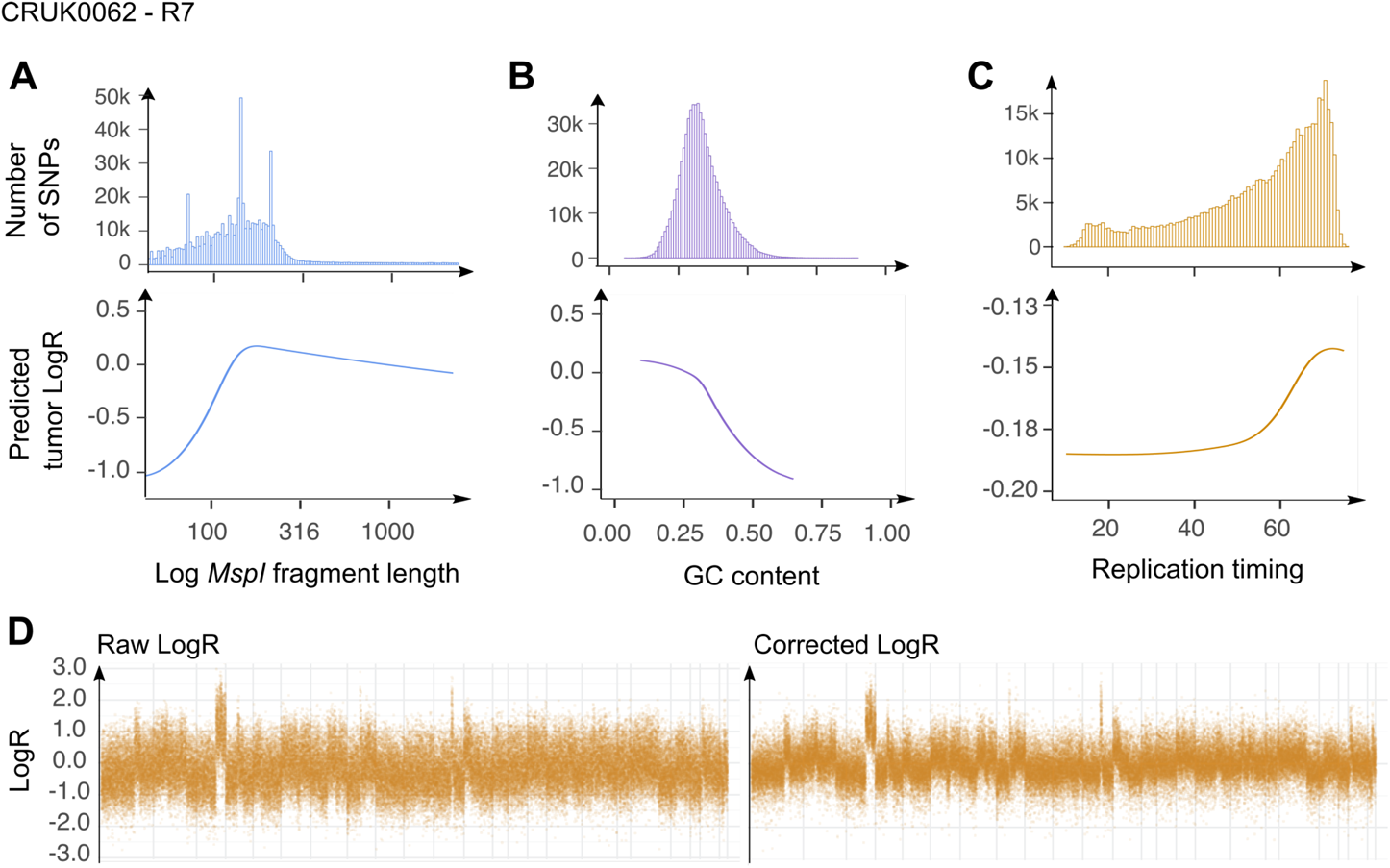
RRBS-derived tumor LogR biases and correction. (A-C) *MspI* fragment length (A), GC content (B) and replication timing (C) affect tumor LogR. (D) A linear combination of three natural splines, modelling the effect of each of the three biases described in (A-C) is used to correct the raw LogR.

**Figure S3.**
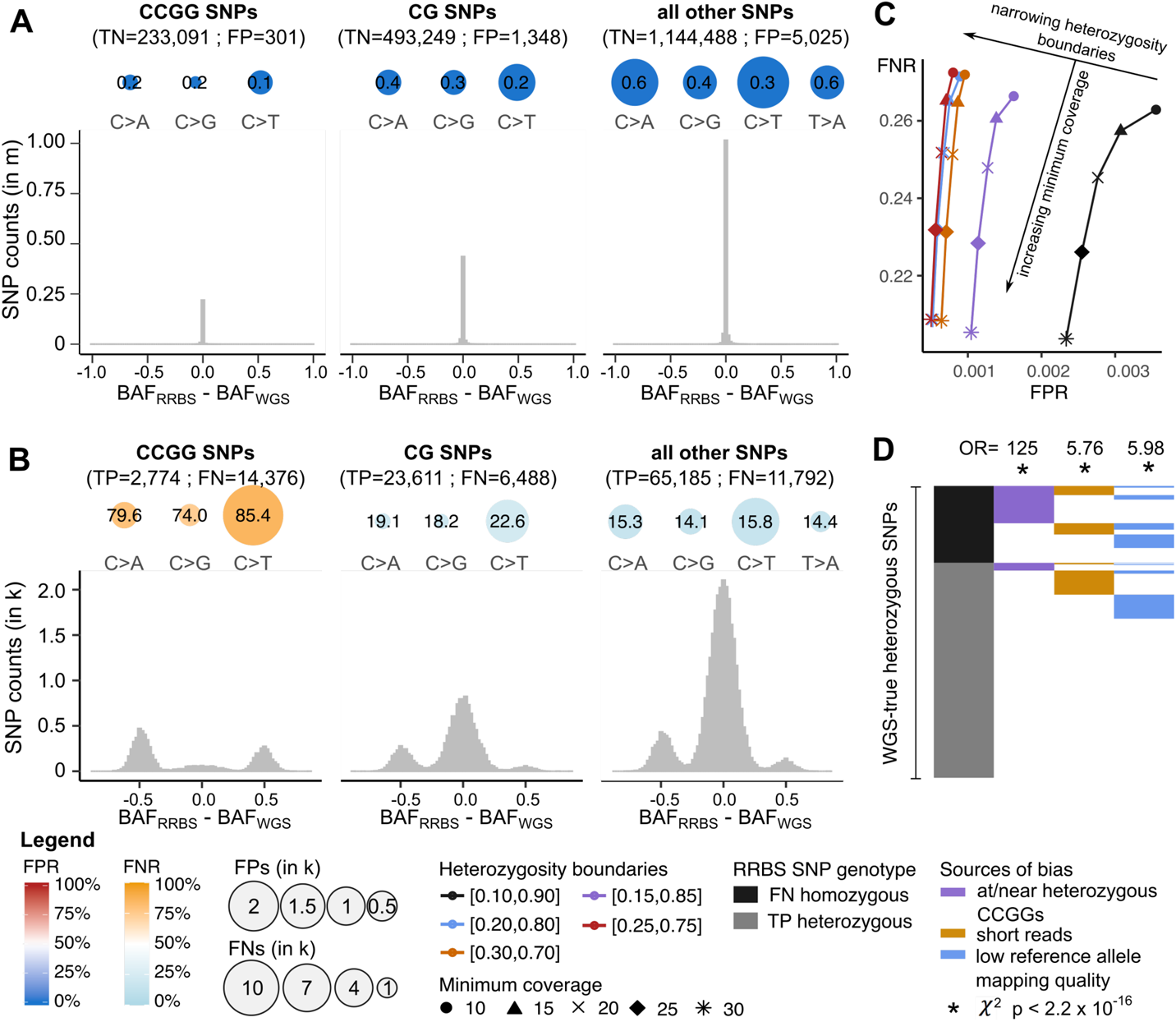
CAMDAC genotyping validation and evaluation of sources of bias affecting heterozygous SNP calling from RRBS data. (A-B) False positive (A) and false negative (B) heterozygous call numbers (scaled radii) and rates (%) stratified by SNP type and sequence context (top) and histograms showing the distribution of CAMDAC BAF estimate errors and noise (bottom). SNPs are considered heterozygous when 0.1 ≤ BAF ≤ 0.9. (C) Performance of heterozygous SNP identification from normal tissue RRBS data at different sequencing coverages and BAF heterozygosity boundaries. (D) Fractions of true positive and false negative heterozygous SNP calls stratified by their location (i) at/or within one read length of a heterozygous CCGG, (ii) covered by only short reads (< 67bp) and (iii) by reads with low reference read mapping quality. ***χ*^2^** tests show enrichment of false positives at each of these three types of regions. The data shown is from the adjacent normal sample of CRUK0062.

**Figure S4.**
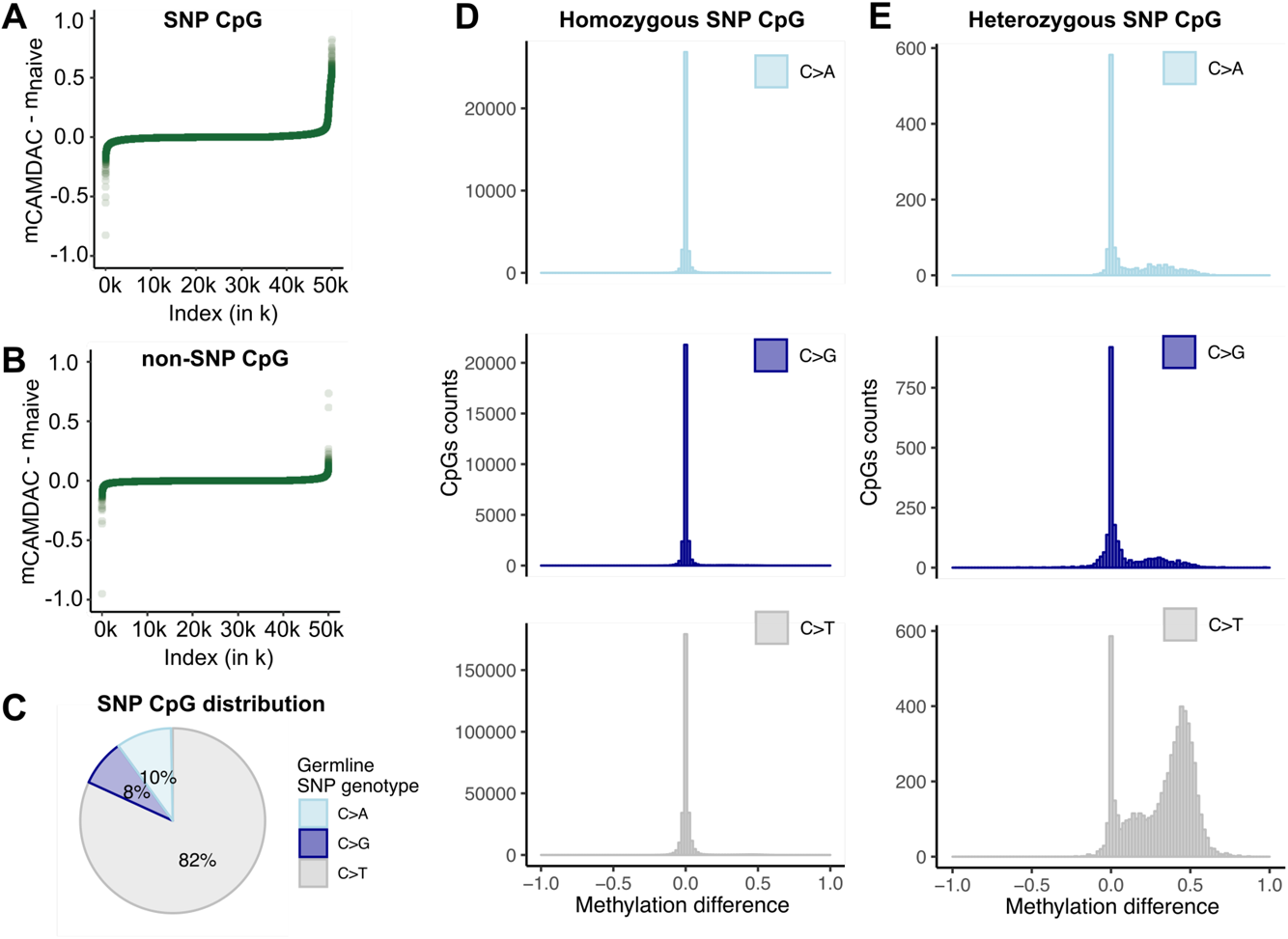
Comparison of CAMDAC and naïve bulk CpG methylation rate. (A-B) Difference between CAMDAC and naïve bulk methylation rate estimates at 50,000 random (A) heterozygous CpGs and (B) non-polymorphic CpGs in a normal sample. (C) Distribution of SNP types at CpGs. (D-E) Difference between CAMDAC and naïve bulk methylation rate values at (D) homozygous *vs.* (E) heterozygous CpG, stratified by type as in (C). All data was taken from the adjacent normal sample of patient CRUK0069, which constitutes a representative example of normal lung methylation in this cohort.

**Figure S5.**
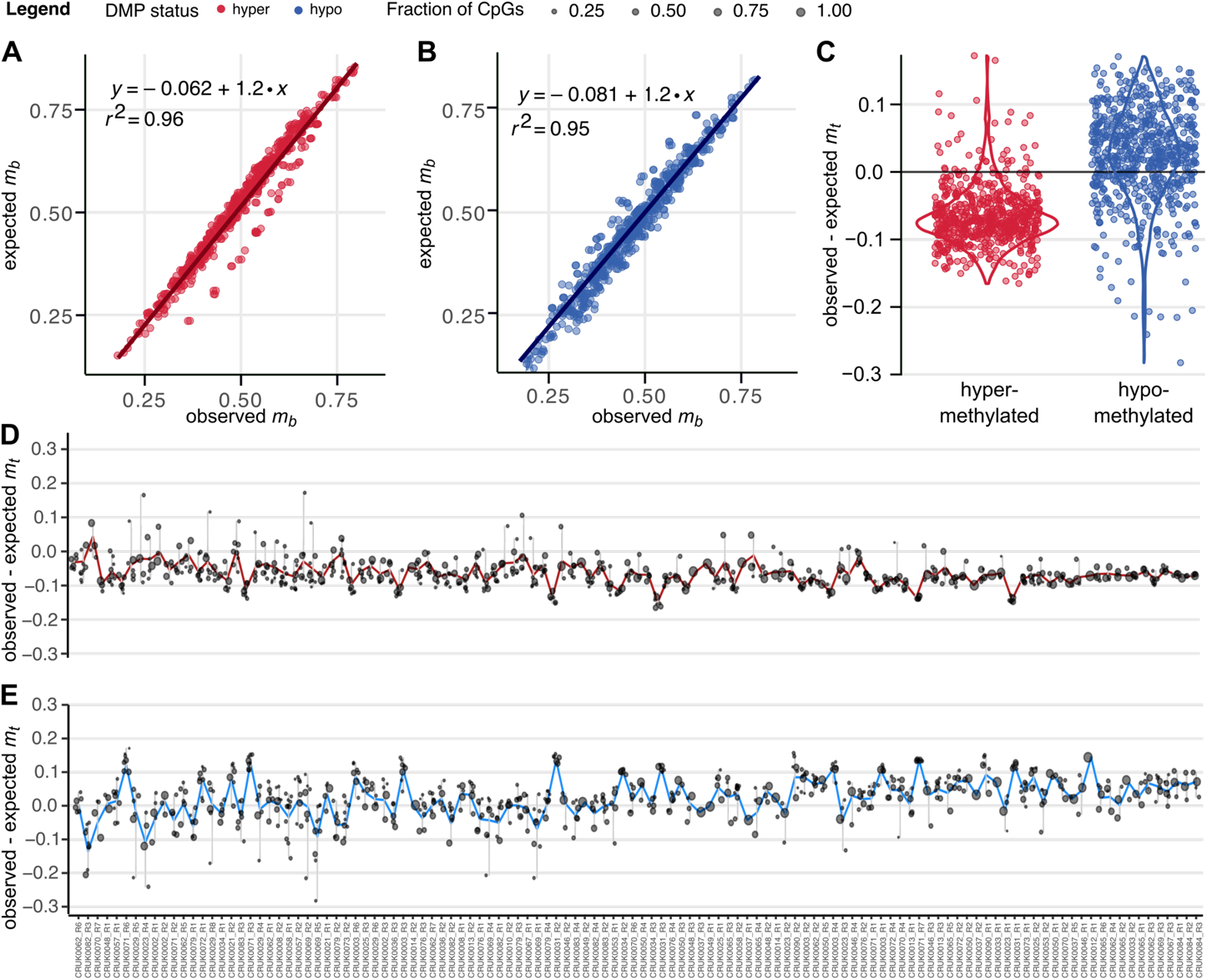
Validation of CAMDAC equations as a function of purity and copy number. (A) Comparison of the predicted methylation rate under the CAMDAC equations as a function of tumor purity and copy number state, and the observed peak position for clonal bi-allelic tumor-normal hypermethylated CpGs (**Methods**). (B) As in A, but for unmethylated CpGs that are methylated in the adjacent normal sample. (C) Observed minus expected CAMDAC pure tumor methylation rate at clonal hypo- (blue) and hypermethylated (red) DMPs across samples (purity) and tumor copy numbers. (D-E) Observed minus expected CAMDAC pure tumor methylation rates at clonal hyper- (D) and hypomethylated (E) CpGs across copy number states and tumor samples.

**Figure S6.**
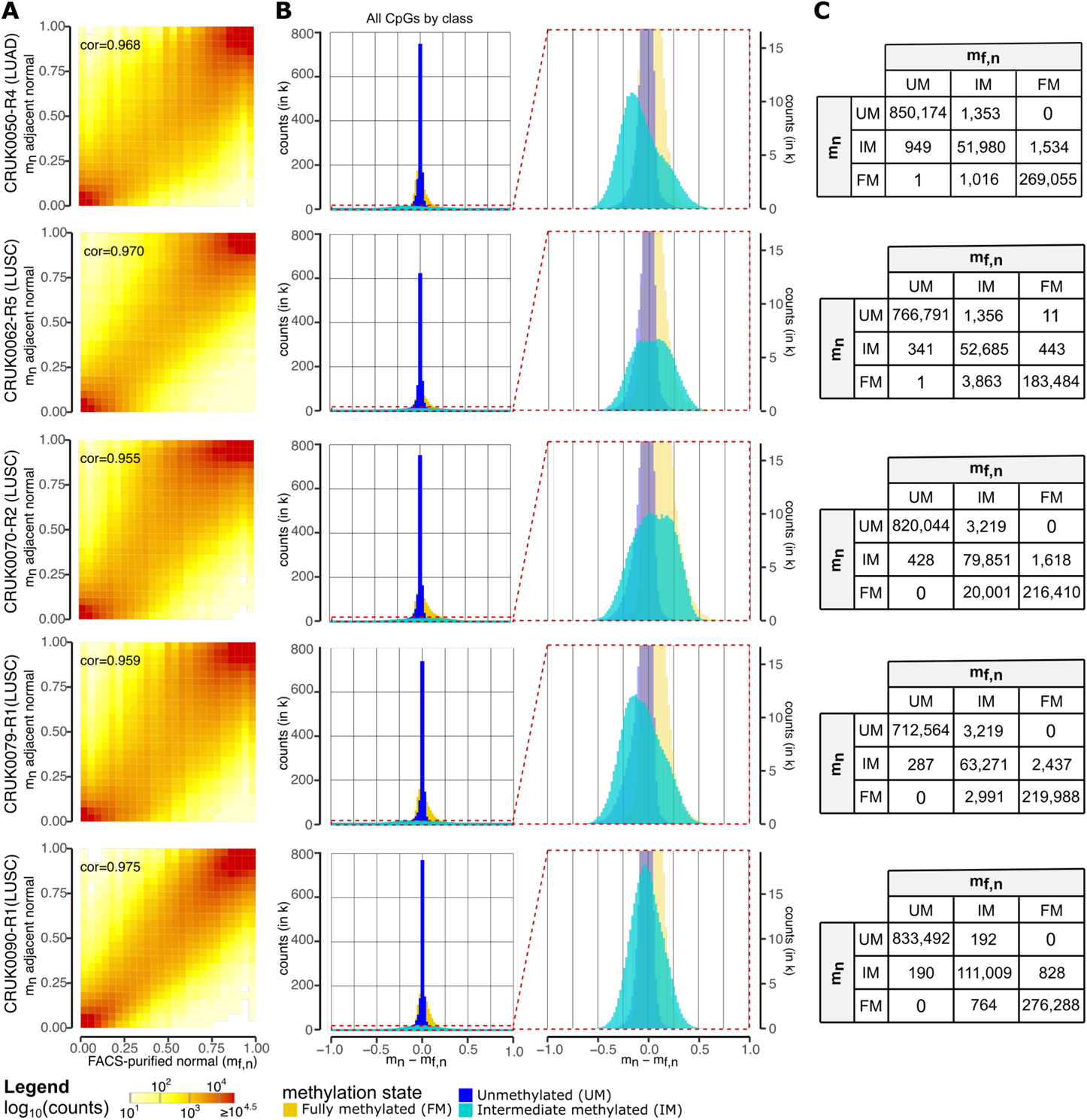
Comparing FACS-purified and adjacent normal methylomes. (A) CpG-wise comparison of the adjacent (*m_t_*) and FACS-purified (*m_f,n_*) normal methylation rates with Pearson correlation. (B) Methylation rate difference histogram for loci classified as confidently unmethylated (UM, HDI^99^ ⊆ [0, 0.2], blue), intermediate (IM, HDI^99^ ⊆ [0.2, 0.8], turquoise) and fully methylated (FM, HDI^99^ ⊆ [0.8, 1], yellow) in at least one sample for a given patient-matched normal pair (first column). Zooming in on the CpGs classed as IM (second column). (C) CpG classification based on *m_t_* and *m_f,n_*.

**Figure S7.**
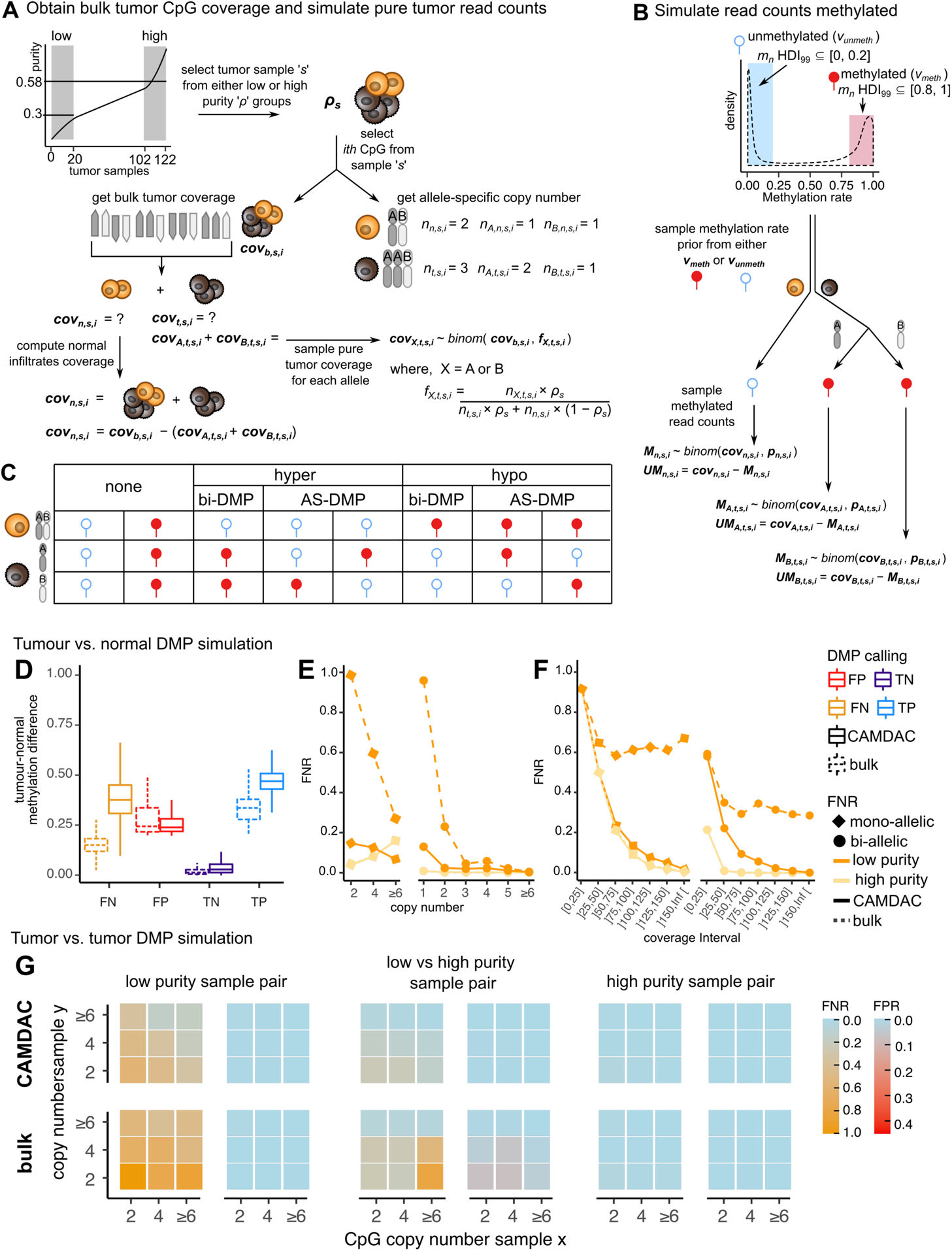
DMP simulation framework and results of tumor-normal and tumor-tumor mono-allelic DMP simulations. (A) Tumors are ranked by purity and the top 20 (high purity; *ρ_s_* ≥ 0.58) and bottom 20 (low purity; *ρ_s_* ≤ 0.3) cases are considered for simulations. Sample *s* with purity *ρ_s_* is randomly chosen from either group, for example, *ρ_s_* = 0.6. Autosomal CpGs are separated into groups based on total copy number *n*_*t,s,i*_ = 1, 2, 3, 4, 5 or ≥6. We then sample the bulk tumor coverage, *cov*_*b,s,i*_, and (allele-specific) tumor copy numbers of CpG *i* with total, major and minor allele copy number, *n*_*t,s,i*_, *n_A,t,s,i_* and *n_B,t,s,i_*, respectively. The matched normal copy number is *n*_*n,s,i*_ = 2. We obtain the pure tumor coverages for alleles A and B, *cov_X,t,s,i_*, where X is either A or B, by sampling from a binomial with the number of trials set to *cov*_*b,s,i*_ and the likelihood of obtaining a tumor read equal to the tumor fraction, *f_X,t,s,i_*, estimated as a function of copy numbers and tumor purity. The matched normal coverage, *cov*_*n,s,i*_, is taken as the difference between the bulk and pure tumor coverages, *cov*_*n,s,i*_ = *cov*_*b,s,i*_ − *cov*_*t,s,i*_. (B) We take confidently unmethylated and methylated loci and collate their methylation rate values into vectors *v*_*unmeth*_ and *v*_*meth*_, respectively. We sample a normal methylation rate prior, *p*_*n,s,i*_ from *v*_*unmeth*_ or *v_meth_*. Leveraging the inferred normal coverage (see A), we sample the counts methylated from a binomial, *M*_*n,s,i*_. The unmethylated reads counts, *UM*_*n,s,i*_, is the difference between the normal coverage and the number of methylated reads, *UM*_*n,s,i*_ = *cov*_*n,s,i*_ − *M*_*n,s,i*_. We then repeat these steps for tumor allele A and B. (C) Possible combinations of normal and allele-specific tumor methylation priors and associated ground truth tumor-normal DMP calls to be compared with CAMDAC DMP calls. (D-F) Results of tumor**-**normal DMP simulations. (D) Mean methylation difference across false negatives (FN), false positives (FP), true negatives (TN) and true positives (TP) at mono-allelic differentially methylated and non-differentially methylated CpGs. (E-F) False negative rates as a function of tumor copy number (E) and coverage (F) for the purified (solid line) and bulk tumor (dashed line) for low *vs.* high tumor purity samples (pale and dark orange lines respectively) at mono- (left panel) and bi-allelic (right panel) epimutations. (G) Results of tumor-tumor DMP simulations. False negative and false positives rates as a function of tumor copy number for low (left panel), low versus high (middle panel) and high purity (right panel) sample pairs at mono-allelic epimutations.

**Figure S8.**
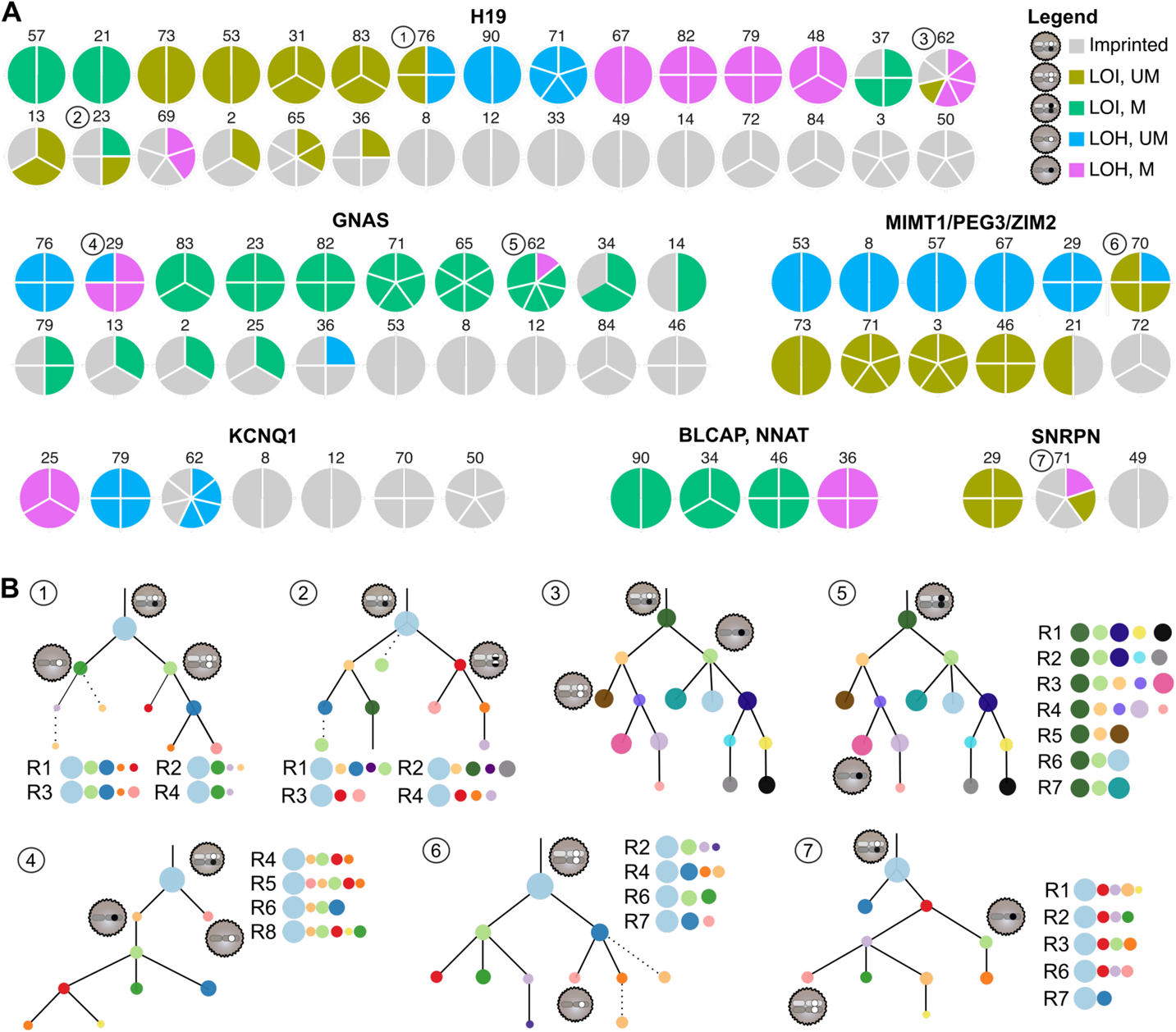
Loss of imprinting ubiquity analysis reveals instances of parallel evolution. (A) Intra-tumor heterogeneity overview of somatic changes at known imprinted loci. (B) Mapping of heterogeneous loss of imprinting events to patient phylogenies.

**Figure S9.**
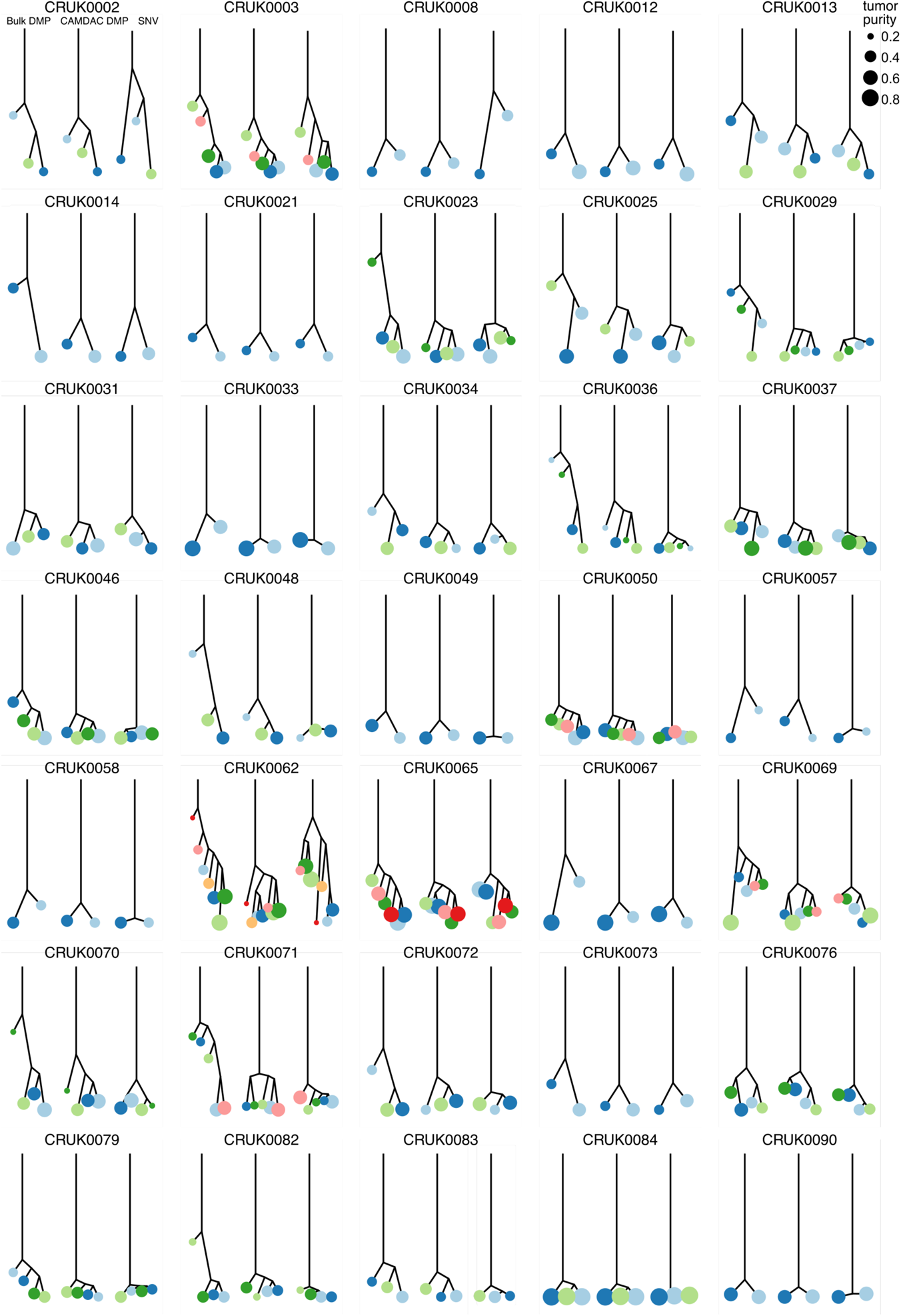
DMP- and SNV-derived sample trees. Phylogenetic sample trees created by calculating the pairwise hamming distance between samples using SNVs from WES or (hypermethylated) tumor-normal DMPs from RRBS, followed by neighbor-joining (**Methods**).

**Figure S10.**
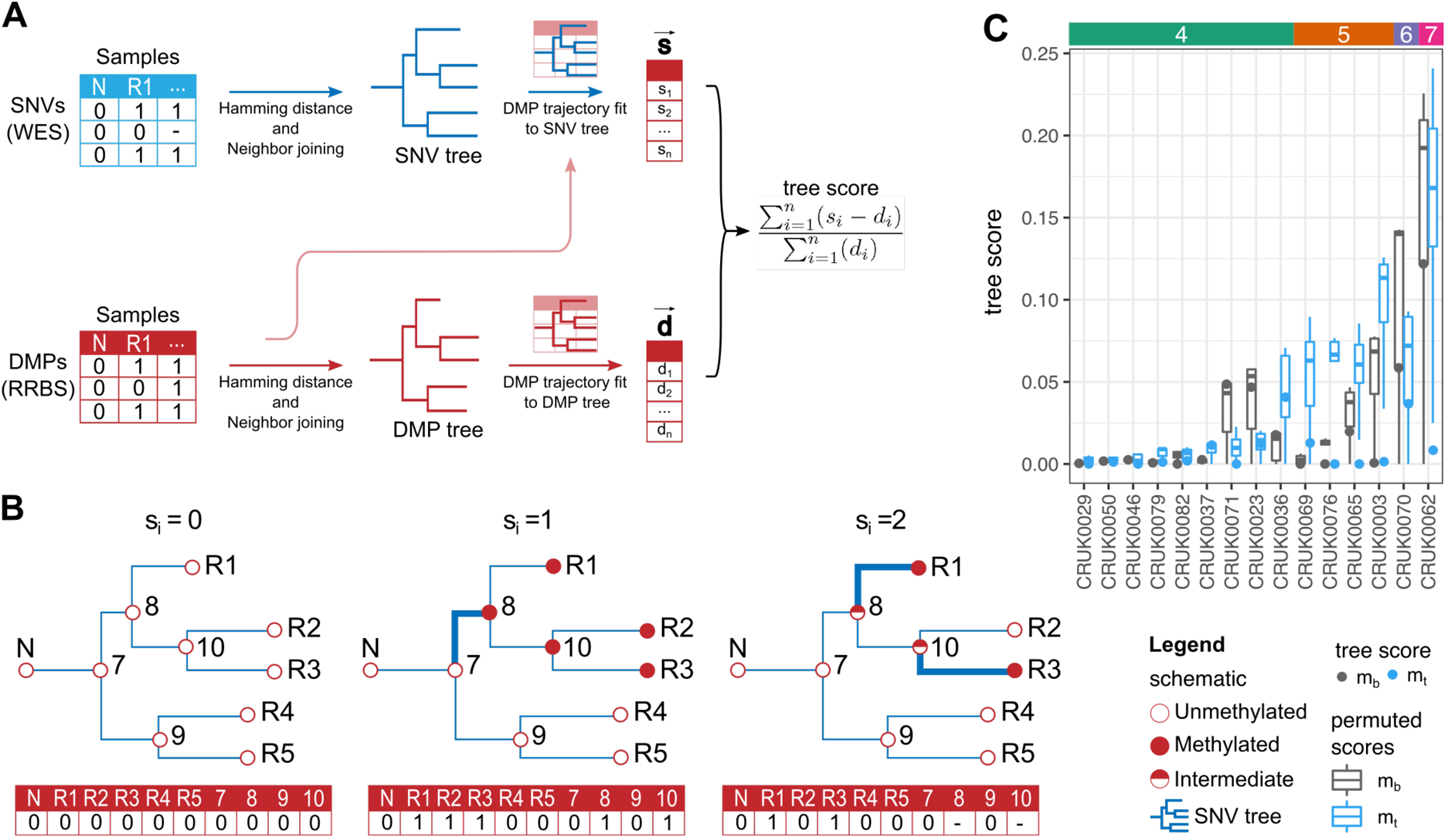
Methylation-SNV tree fit scores. (A) Scoring concordance between phylogenies derived from aberrant methylation and somatic sequence alterations. We leveraged binarized (hypermethylated) tumor-normal DMP data and, based on the ancestral states given by either the SNV (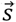) or DMP (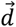) multi-sample tree topologies, counted the number of state changes per DMP locus from the normal root to all leaves (methylated to unmethylated or *vice versa*). Counts were combined into a single score by taking the total absolute difference in the number of ancestral DMP state changes based on the SNV (*s_i_*) and DMP tree topologies (*d_i_*, 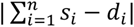) as a fraction of all events given by the DMP tree (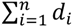). Here, a score of 0 indicates a perfect topology match, with the score increasing with the number of inconsistent per-DMP evolutionary trajectories between the two trees. Hence, the score indicates how closely the methylation events support sample relationships inferred from somatic sequence alterations. (B) A schematic of how ancestral state changes are calculated given the SNV tree by counting edges that switch from methylated to unmethylated (bold). Internal node states are inferred using ancestral reconstruction and, for all calculations, nodes with intermediate methylation are replaced with the state of the first ancestor without intermediate methylation. (C) Methylation-SNV fit scores calculated for DMPs using *m_b_* and CAMDAC *m_t_*. Boxplots show the distribution of scores for permutated topologies in place of the SNV tree, as used for empirical p-value calculations. A color bar indicates the number of tumor regions per patient (excluding the patient-matched normal).

## METHODS

### TRACERx methylation study

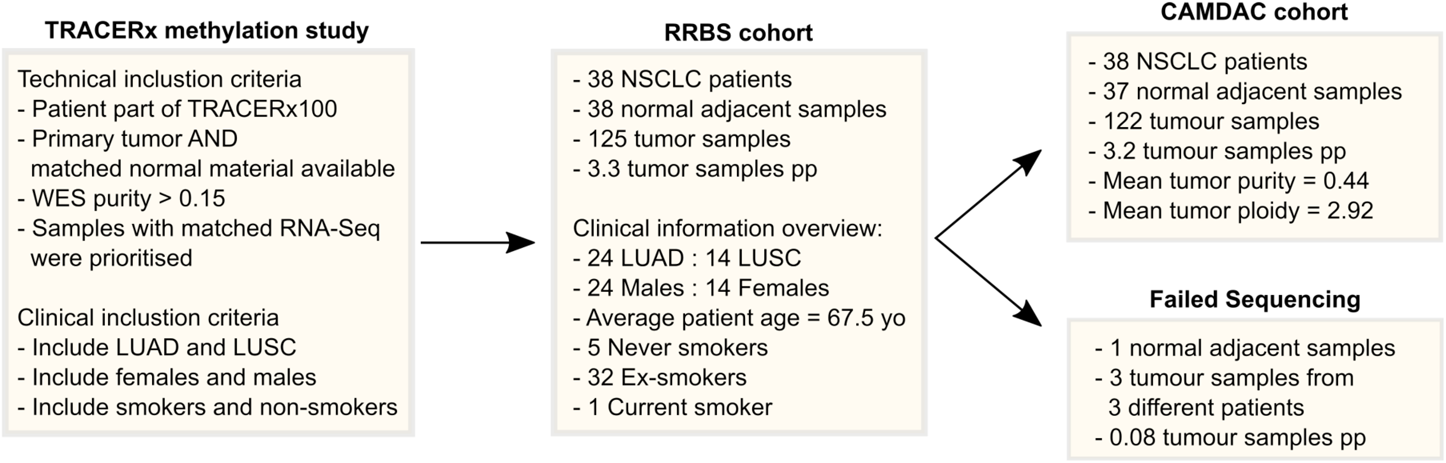

Samples from the first 100 patients of the TRACERx lung cancer study were selected for reduced representation bisulfite sequencing (RRBS). Patients with sufficient material remaining from 2 or more tumor regions and the adjacent matched normal were considered for bisulfite sequencing. Samples with purity below 15% were discarded with the exception of CRUK0062-R6 included for comparison with the other 6 tumor regions sampled for this patient. Patients with tumor samples of high purity were prioritized as well as those with matched RNA-Seq data (Rosenthal et al., 2019). We also chose samples from both lung adenocarcinoma (LUAD) and squamous cell carcinoma (LUSC) subtypes, genders and smoking status.

The TRACERx WES, WGS and RRBS data generated, used or analyzed in this study are not publicly available and restrictions apply to the availability of these data. These data are available through the Cancer Research UK & University College London Cancer Trials Centre for academic non-commercial research purposes upon reasonable request, and subject to review of a project proposal that will be evaluated by a TRACERx data access committee, entering into an appropriate data access agreement and any applicable ethical approvals.

### Reduced representation bisulfite sequencing

RRBS was obtained for roughly a third of the NSCLC patients from the TRACERx 100 cohort (122/327 tumor regions from 38/100 patients, 37 with matched normal). The NuGEN Ovation RRBS Methyl-Seq System, adapted by the manufacturer for automation on Agilent Bravo liquid handling robot, was used to generate sequencing libraries by enzymatically digesting 100 ng of gDNA using *MspI*, which recognizes 5′-CCGG-3′ sequences and cleaves the phosphodiester bonds upstream of CpG di-nucleotide, leaving a 2bp overhang suitable for adaptor ligation and then a final end repair step. Generated libraries were bisulfite converted using Qiagen’s EpiTect Fast DNA Bisulfite Kit. Converted libraries were amplified by PCR using 12 cycles and purified using Agencourt® RNAClean® XP magnetic beads. Purified libraries were quantified by Qubit dsDNA HS Assay (Invitrogen) and quality was evaluated using Agilent Bioanalyzer High Sensitivity DNA Assay (Agilent Technologies). Eight samples were multiplexed per flow cell and sequenced on HiSeq2500 system using HiSeq SBS Kit v4 in paired-end 100bp runs for CRUK0062 and single end 100 bp runs for the others yielding an average of 150M raw sequencing reads per sample.

Sequencing outputs were converted to FASTQ files using Illumina’s bcl2fastq Conversion Software, quality checked with FastQC v0.11.2 (Babraham Institute, https://www.bioinformatics.babraham.ac.uk/projects/fastqc/) and adapter sequences and diversity bases were trimmed with Trim Galore! version v0.3.7 (Babraham Institute, https://www.bioinformatics.babraham.ac.uk/projects/trim_galore/), a wrapper around Cutadapt, (Martin, 2011) and NuGEN’s trimRRBSdiversityAdaptCustomers.py v1.0 (https://github.com/nugen technologies/NuMetRRBS). Reads are aligned to the UCSC hg19 reference assembly using Bismark v0.14.4 (Krueger and Andrews, 2011) and deduplication was carried out using NuDup, leveraging NuGEN’s molecular tagging technology (https://github.com/nugentechno logies/nudup). Binary alignment map (BAM) files were then sorted and indexed using SAMtools v1.2 (https://github.com/ samtools/samtools/releases/tag/1.2).

On average 1.88 × 10^8^ reads per sample remained post-processing and alignment (**Table S2**), resulting in an average of 4.5 million CpGs being supported by at least one read in any one sample.

### Whole genome sequencing

Whole genome sequencing was performed on 8 samples from 3 patients included in the TRACERx100 cohort. Simultaneous extraction of DNA and RNA was performed using the AllPrep DNA/RNA Mini Kit (Qiagen, UK). Frozen samples were transferred onto cold petri dishes on dry ice and were manually dissected into 20-30mg pieces. Immediately prior to extraction, the freshly dissected tissue was transferred directly into homogenization tubes containing RLT plus lysis buffer. Homogenization of tissues was carried out using a TissueRuptor II probe or using a bead method and by passing the lysate through a QIAshredder column (Qiagen, UK). The extracted DNA was eluted with 200 µl of EB (no EDTA) buffer and RNA was eluted with 200µl of nuclease-free water and stored immediately at −80°C. The DNA and RNA samples were quantified using a Qubit™ 3.0 Fluorometer (Lifetechnology, UK) and TapeStation system (Agilent, UK) respectively. The integrity of the DNA/RNA was assessed using the Agilent TapeStation system.

Whole genome sequencing experiments were conducted by Edinburgh Genomics. Samples were sequencing on Illumina HiSeq X in paired-end 100bp runs. FASTQ outputs sequencing reads were quality checked with FastQC v0.11.5 (Babraham Institute, https://www.babraham.ac.uk/) and then aligned to the UCSC hg19 reference assembly (including unknown contigs) obtained from GATK bundle 2.8 using bwa-mem v0.7.15 (http://bio-bwa.sourceforge.net). Picard tools v1.107 was used to clean, sort, de-duplicate and merge files from the same patient region and to remove duplicate reads (http://broadinstitute.github.io/picard). Binary alignment map (.bam) files were sorted and indexed with SAMtools v1.3.1 (http://github.com/samtools/) setting both the minimum base and mapping quality to 20.

### Copy-number aware methylation deconvolution analysis of cancers (CAMDAC)

As depicted below, CAMDAC requires RRBS data prepared from bulk tumor and matched adjacent normal samples. CAMDAC is only compatible with human (directional) RRBS data. The input must be quality and adapter trimmed with PCR duplicates removed and subsequently aligned to hg19 (hg38, GRCH37 and GRHCH38 formats also compatible). The tumor and matched normal data must be provided in .bam file format. The files must be sorted and indexed using SAMtools (http://github.com/samtools/). The input tumor and matched normal sequencing data is used to compute tumor purity and allele-specific copy numbers as well as bulk tumor and normal methylation rates. From this, we extract purified tumor methylation rates and carry out tumor-normal and tumor-tumor differential methylation analysis.

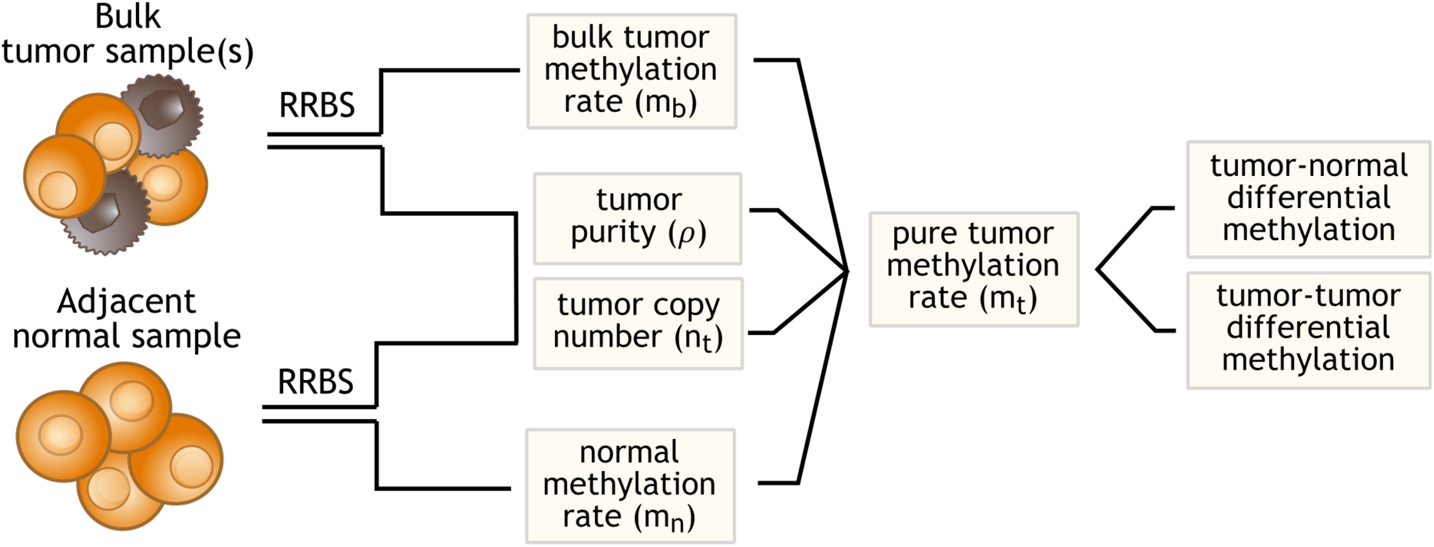

We created a reference RRBS genome listing all genomic regions supported by at least 5 reads in at least one of the 37 germlines from this study. Genomic regions which fail to reach this criterion are unlikely to achieve minimum depth thresholds required for copy number and differential methylation analysis and can be excluded, ultimately to speed up computation.

### Tumor copy number profiling from RRBS data

First, we used CAMDAC get_allele_counts() command (https://github.com/VanLoo-lab/CAMDAC, build=“hg19”, mq=0, n_cores=12) to compile base counts for SNP loci described by the 1000 Genomes project (1000g) (Auton et al., 2015) which overlap with the above-mentioned list of genomic regions. By default, no mapping quality threshold is set to avoid creating a bias against the alternate allele at SNPs. Allele counts are obtained for each of these SNP to calculate the BAF and LogR.

The LogR value at SNP *i* represents the base-2 logarithm of the ratio of the coverage in the tumor (*cov_t,i_*) to that of the normal at that SNP (*cov_n,i_*), divided by the average ratio. This is easily computed from RRBS data, as shown in *Eq(4)*.

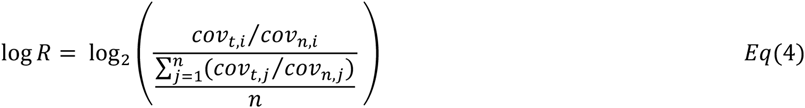

Normalization with the adjacent normal SNP coverage, removes germline copy number variants. Next, we correct these germline normalized LogR values for fragment length, GC content and replication timing biases (**Figure S2**). This is a necessary process post-normalization as technical and biological biases affecting the sequencing output can vary between the normal and bulk tumor data. In the first instance, we reconstruct the underlying *MspI* fragment distribution from the sequencing output for each patient. Assuming, and confirming, complete enzymatic digestion, *MspI* fragment ends can be found at CCGG motifs, taking into account SNP forming/destroying cleavage sites. A CCGG is confirmed when supported with a 5’ read starting with one of the following trinucleotides: CGG(+), TGG(+), CCG(−) or CCA(−). Having identified the *MspI* fragments, we can readily annotate the fragment length. Then, we compute the replication timing estimate at every 1000g SNPs leveraging Repli-seq data from 15 cell lines produced by the ENCODE project (Dataset GEO Accession: GSE34399), generating 15 reference profiles. The profile which best fits the LogR data in a given tumor sample is then used to calculate the replication timing for each *MspI* fragment in that sample. To compute the GC content of all fragments, we leverage information from the bulk tumor reads, adjusting the reference GC content for methylation rate and bisulfite conversion. Where the fragments are longer than 2x read length, we assume that all reference cytosines are converted to thymines unless they are at CpG where we assume a probability equal to the mean of all informative CpGs loci in the fragment. Finally, we fit the observed LogR to a linear combination of the natural splines (df = 5) of *MspI* fragment length, replication timing and GC content. The model residuals provide corrected LogR values.

The BAF value at the i^th^ 1000g SNP is the ratio of the alternate allele counts to the total counts at the ith SNP locus, 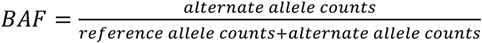. The strand-specific BAF rules for every SNP type are shown in **Figure 2D**. Here, we make the reasonable assumption that all reference cytosines are converted into thymines (under-conversion rate <0.5%). Note that where a variant makes up more than 5% of the total allele counts at a given SNP, it is excluded from downstream analyses in order to ensure that SNVs or mis-alignments do not affect BAF estimates and subsequent copy number profiling.

Where allelic imbalance is such that two distinct band are generated, CAMDAC can assign heterozygous SNPs (0.3 ≤ BAF ≤ 0.7) to either the gained or lost alleles. SNP phasing information can then be shared between tumor samples from the same patient. This can be used to identify mirrored subclonal allelic imbalance (Jamal-Hanjani et al., 2017), but most importantly, it allows us to rescue signal in tumors where there was no clear separation before haplotyping.

With partially phased BAF and corrected logR estimates in hand, CAMDAC then uses ASCAT (Van Loo et al., 2010) piece-wise constant segmentation (penalty = 200) leveraging germline heterozygous SNPs (0.3 ≤ BAF ≤ 0.7) and copy number fitting functions (gamma = 1) to obtain allele-specific copy number profiles and purity estimates for each tumor region. Note that we set the minimum germline coverage for SNP inclusion to 10 whilst one read is deemed sufficient in the tumor. To remove outliers and further reduce noise without introducing a bias against homozygous deletions, we remove singletons with coverage below 10 in the tumor if and only if their nearest neighbors within a 10kb moving window all pass the above threshold.

There were 6 patients which showed large intra-tumor differences in their CAMDAC ploidy estimates. In each of those cases, we looked for evidence supporting an alternative ploidy solution. If such a solution existed and was better suited the overall tumor ploidy profile, the ASCAT copy number fitting step was re-run forcing the solution to the ploidy towards a diploid or tetraploid solution (see MINPLOIDY and MAXPLOIDY variables, from the runASCAT() function in ascat.R script; available from https://github.com/VanLoo-lab/ascat). We refit 7 tumor copy number profiles in total. This is an improvement from the 17 samples which were manually curated and re-run in the matched exome sequencing data (Jamal-Hanjani et al., 2017).

### Copy number gains and losses summary

We calculated the fraction of gains and losses in LUAD and LUSC samples using copy number alterations called with CAMDAC. The genome was partitioned into 10MB bins, each of which were classified per sample: a “gain” if the major allele is above the baseline ploidy and a “loss” if the minor allele falls below the baseline ploidy and the total tumor copy number is lower than 5. A given genomic bin can therefore be classified as both a gain and loss. The baseline ploidy is defined as diploid or tetraploid dependent on ASCAT solution. Fractions of each class were calculated across the cohort, with 99.99% (530 / 536) of bins with coverage in all samples. For TCGA samples, copy number alterations were obtained from SNP array data with ASCAT. TCGA gains and losses were obtained as above and compared with those of the TRACERx cohort by taking the average fractions at the corresponding 10MB bins. Where TCGA SNP array data did not cover RRBS regions (bins on chromosome 1 and 9), the value of the previous segment is taken as an estimate.

### Comparing RRBS- and WGS-derived genotypes

First, we compiled allele counts at all 1000g SNPs with coverage in the matched RRBS data using alleleCount v4.0.0 (http://cancerit.github.io/alleleCount/). We compared CAMDAC RRBS-derived genotypes with the output from standard alleleCount v4.0.0 pipeline on WGS data for the 3 germline samples of patients CRUK0031, CRUK0062 and CRUK0069. Positive calls are heterozygous SNPs (0.1 ≤ BAF ≤ 0.9) and WGS is considered to provide ground truth with a minimum SNP read depth of 10 required on both platforms for inclusion. We compute the average false positive rate (FPR) as the sum of all false positives over the sum of false positives and true negatives across samples. Similarly, the mean false negative rate (FNR) is calculated as the sum of all false negatives divided by the sum of false negatives and true positives. The FPR allows us to evaluate allele-specific biases introduced by the approach, while the FNR assesses the influence of methylation status on BAF calculation, as well as biases introduced during the RRBS protocol. For all SNP types, we consider three different contexts: (i) SNPs at CCGG, the recognition motif of the *MspI* restriction enzyme used during library preparation, (ii) SNPs at CpGs (excluding CCGG motifs), and (iii) all other. A chi-square analysis is performed to test whether or not the FNR is context dependent. Next, we recompute FNR and FPR aggregating FP, TP, FN and TN across all samples for a range of heterozygosity boundaries and minimum coverage thresholds (**Figure S4A**). Next, we measure the number of false negatives due to (i) allele-specific *Msp1* fragments, (ii) additional reference mismatches from neighboring SNPs, and (iii) short fragments yielding invalid alignments given 2 or less reference mismatches (**Figure S4B**).

The mean ASCAT.m FPR across all SNPs and samples was 0.3% and was always below 2%. The average FNR was 25% and was highly context dependent (*χ*^2^ test, p-val < 2.2 × 10^−16^, FNR_CCGG_ = 83%, FNR_CG_ = 20%, FNR_other_ = 15%). Higher sequencing coverage results in only modest improvements in FNR: by requiring a minimum SNP coverage of 30 instead of 10 reads, the average FNR dropped to 20% (**Figure S4A**). As expected, the majority of heterozygous SNPs perturbing or creating an *MspI* recognition motif are missed (49% of all false negatives, **Figure S4B**). These polymorphisms result in allele-specific fragments during RRBS library preparation that skew allelic coverage. Heterozygous SNPs on the same *MspI* fragment as a heterozygous CCGG are similarly affected. Finally, within the false negative calls, we see a bias towards SNPs erroneously called homozygous reference (72%) compared to homozygous alternate (28%), and this effect is present in all three contexts, although less pronounced at CCGGs. This points towards an alignment bias, possibly explained by limited mappability of short *MspI* fragments with alternate alleles. Supporting this theory, SNPs with lower reference allele read mapping quality scores were more likely to yield homozygous reference false negatives (Wilcoxon test, p < 2.2 × 10^−16^), especially outside CCGG context.

### Tumor copy-number profiling from whole genome sequencing data

First, we compiled allele counts at all 1000g SNPs with coverage in the matched RRBS data using alleleCount v4.0.0 (http://cancerit.github.io/alleleCount/). We then we ran ASCAT GC and replication timing LogR correction steps, fed the output into ASCAT piece-wise constant segmentation (penalty = 200) and obtained copy number and purity (gamma = 1). Comparing WGS and RRBS-derived profiles, even with the higher FNR on SNP calling in RRBS data, CAMDAC BAF estimates enable ASCAT to correctly identify separation of the BAF bands on several occasions where the WGS BAF does not seem to provide sufficient power. Likewise, comparison of the LogR tracks indicates that RRBS protocol-induced biases in the coverage are adequately modelled and removed by CAMDAC. Unsurprisingly, final allele-specific copy number segments are virtually indistinguishable between the two data types.

### Methylation rate calculation from RRBS data

The bulk tumor and matched normal methylation rate is readily computed by taking the ratio of methylated CpG read counts to the sum of methylated and unmethylated read counts, 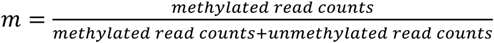. CAMDAC uses strand-specific dinucleotide counts to distinguish between methylated and unmethylated CpGs, 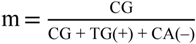. At CpG>TpG (TpG>CpG) and CpG> CpA (CpA> CpG) SNPs, only reads from the bottom strand and top strand, respectively, contribute to CAMDAC methylation rate estimates. In addition, only the CG-forming allele contributes to the methylation rate at polymorphic CpGs. This enables the methylation rate at a heterozygous CpG to vary between 0 and 1, rather than between 0 and 0.5 in a diploid sample and ensures further independence between methylation rate and copy number estimates.

We compiled bulk tumor and normal methylation rates for all CpGs which fell within the above-mentioned reference RRBS genomic regions list. For each patient, we discarded all CpGs that failed to reach a minimum coverage of 10 in the matched normal RRBS data. CpGs that had less than 3 reads in a given tumor sample were also filtered out from that sample.

### CAMDAC purified tumor methylation rates from RRBS data

Bulk tumor methylation rate (*m_b_*) could be expressed as a function of the methylation rate in the tumor cells (*m_t_*) and contaminating normal cells (*m_t_*), scaled for the purity of the sample (*ρ*) and the local copy number state (i.e. *n*_n_ = 2 in the normal and *n*_t_ in the tumor):

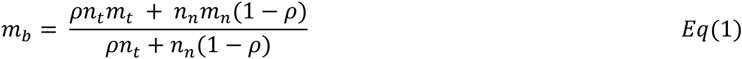

Taking estimates of the total copy number at each CpG and tumor purity from the RRBS copy number profile and assuming the matched normal is a reasonable proxy for the contaminating normal, we can calculate the deconvolved tumor methylation rate *m_t_*:

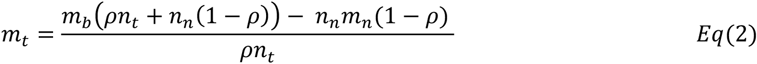

An important distinction is made at heterozygous polymorphic CpGs, where only one copy in the diploid normal contains methylation information (i.e. *n*_&_ = 1). Similarly, if the SNP is CG-destroying and its BAF is smaller than 0.5 or likewise if the SNP is CG-forming and BAF is above 0.5 the major copy informs the methylation (i.e. *n*_t_ = *n_major_*). *Vice versa*, if the tumor BAF is larger than 0.5 and the SNP is CG-destroying or the BAF lower than 0.5 and the SNP is CG-forming, the minor allele only informs the methylation rate (i.e. *n_t_* = *n_minor_*).

Finally, *m_t_* must be between 0 and 1. However, due to both technical and biological noise, values can fall outside of these boundaries. For downstream analyses, we set negative methylation rates up to 0 and values sitting above the upper boundary are rounded down to 1. In virtually all cases, the *m_t_* 99% HDI overlaps with allowed values.

### Nuclei extraction and FACS analysis

We followed the ‘Frankenstein’ protocol (Martelotto, 2020) to extract nuclei from seven fresh frozen tissue samples taken from five different patients: CRUK0050-R4, CRUK0062-R5, CRUK0070-R2 and R4, CRUK0079-R1 and R4 and CRUK0090-R1. Samples were taken from different tissue cuts of the same sampled tumor region as the matched WES and RRBS samples. The bulk tumor tissue samples were minced and put into a lysis buffer. The lysate was homogenized with a pestle and the mixture was then passed through a 70μm filtered. The isolated nuclei mixture was washed and suspended into a buffer containing propidium iodide (PI) stain (70 μg/mL PI, 1% BSA, 1×PBS) and passed through a smaller 35μm filter.

The stained nuclei were then passed through the FACS machine. Nuclei were separated according to the side scatter, a measure of cell morphology and the PI chromophore intensity, representing the quantity of chromatin in each nucleus. We first sorted a small number of nuclei to adjust the sorting parameters to ensure that debris was removed. We also discarded superfluous ploidy populations, including nuclei having originated from replicating diploid or aneuploid cells. We then collected nuclei from both the major aneuploid population, *popA*, and the diploid population, *popD*, into separate tubes each with 200 μL PBS and 2% FCS. We counted between 100-300 thousand events per population. DNA extraction for each sample and nuclei subpopulation was performed using the Zymo Research Quick DNA-microprep plus kit (D4074) following the indications from manufacturer. The DNA samples were finally eluted in 12μL of elution buffer.

### RRBS of FACS-purified samples

The NuGEN Ovation RRBS Methyl-Seq System protocol was followed for library preparation and for subsequent bisulfite sequencing, as above-described for the bulk tumor RRBS data, but without any automation.

Libraries were prepared by enzymatically digesting ∼100ng of gDNA with *MspI*. Qiagen’s EpiTect Fast DNA Bisulfite Kit was used for bisulfite conversion of the resulting DNA fragments. Bisulfite converted libraries were then amplified by PCR using 12 cycles and purified using Agencourt® RNAClean® XP magnetic beads. Library quantification was performed by Qubit dsDNA HS Assay (Invitrogen) and quality control was carried out using Agilent Bioanalyzer High Sensitivity DNA Assay (Agilent Technologies). In cases with two samples from the same patient, both *popA* and only one *popA* were made into libraries and sequenced to save on costs. We therefore sequenced 7 tumor aneuploid and 5 (presumably) normal diploid populations.

RRBS was performed by at the Francis Crick Institute sequencing facility. The 12 samples were multiplexed across 4 lanes on HiSeq 4000 using the HiSeq® 3000/4000 SBS Kit. As for the bulk tumor samples, 100bp SE and 10bp reads were generated for the NuGEN RRBS library insert and unique molecular identifiers, respectively. We aimed to sequence 120,000,000 reads per sample.

Sequencing reads were QC’ed, adapter trimmed, aligned to hg19, PCR-deduplicated and output binary alignment map (BAM) files were sorted and indexed following the same procedure as described for the TRACERx bulk tumor RRBS dataset (Methods, section 2.4.2.1).

On average, 109,864,415 raw sequencing reads were obtained per samples. Mapping efficiency averaged around 70.5%, as expected for bisulfite sequencing data aligned with Bismark. However, samples had very high duplication rates (average 57.18%), leaving only 31,533,497 reads per sample post-processing. This is potentially due to lower DNA inputs and combined bisulfite- and FACS-driven DNA degradation. This effect was not observed on the previous cohort generated using the same protocol, but without FACS.

### SNV calling from whole-exome and -genome sequencing data

SNV calls from TRACERx 100 patients are readily obtained Jamal-Hanjani *et al*. (Jamal-Hanjani et al., 2017). The same protocol is followed to identify SNVs from newly generated WGS data and is detailed below. In short, SAMtools mpileup (v1.3.1) was used to locate alternate alleles in tumor and germline samples (minimum base quality ≥ 20 and mapping quality ≥ 20). The output was processed with VarScan2 v2.3.6 processSomatic and used to identify somatic variants (VAF ≥ 0.01, purity = 0.5), filtering out germline SNPs using the matched germline samples. The resulting single nucleotide variant (SNV) calls were filtered for false positives using Varscan2’s associated fpfilter.pl script, initially with default settings then repeated again with min-var-frac = 0.02, having first run the data through bam-readcount v0.5.1 (https://github.com/genome/bam-readcount). SNVs were also identified using MuTect v1.1.4, filtered for passing SNVs (“PASS”) and annotated using the GATK bundle 2.8. SNVs were deemed true positives if (A) VAF ≥ 2% and the SNV was called by both VarScan2, with a somatic p-value ≥ 0.01, and MuTect, (B) VAF ≥ 5% and the mutation was only called in VarScan2, again with a somatic p-value ≤ 0.01. A minimum total and mutant allele coverage of 30 and 5 reads was required for inclusion. The number of reads supporting the variant in the germline data must to be < 5 and the VAF ≤ 1%. SNVs were also removed if they were found to match a germline polymorphisms with population frequency > 1% across the TRACERx 100 cohort. SNVs overlapping with a blacklist of genomic regions including reports by the ENCODE project (both DAC and Duke list), simple repeats, segmental duplications and microsatellite regions were obtained from the UCSC Genome Table Browser.

By sharing mutations calls between samples from the same patient, and reassessing mutant reads at each position, we can reduce the possibility of over-representing the mutational heterogeneity. Where a somatic variant was not called ubiquitously across tumor regions, but was called in one or more sample, reads were re-extracted from the original alignment file using bam-readcount v0.5.1 (https://github.com/genome/bam-readcount). In these instances, minimum VAF requirements were decreased to VAF ≥ 1%, thereby rescuing low frequency variants that would otherwise have been missed.

### SNV-phased methylation rate estimates

Leveraging SNV calls from newly obtained WGS data and previously published WES (Jamal-Hanjani et al., 2017), we phased CpG methylation to all SNVs, excluding loci with VAF ≤ 0.1 in a tumor sample or VAF > 0 in the patient-matched adjacent normal tissue. Allele-specific methylation counts were compiled for all reads that could be phased to exactly one SNV. Phased methylation rates were obtained for 32,874 CpGs and 6,529 SNVs across samples (14,514 and 2,984 unique CpGs and SNVs respectively across patients). The VAF was derived from the mutant (mut) and wild type (WT) reads counts: 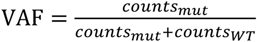. RRBS-derived VAF estimates were compared with those obtained from the WES/WGS (Pearson correlation = 0.86).

The mutation copy number, *n_mut_*, was then computed as a function of the variant allele frequency, tumor purity and copy number: 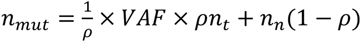. The wild type allele copy number, *n*_WT_, is obtained by subtracting *n_mut_* from *n*_t_: *n_WT_* = *n*_t_ − *n_mut_*. The mutant allele methylation rate, *m_mut_*, is extracted by taking the counts methylated (*m_mut_*) and unmethylated (*UM_mut_*) divided by all counts phased to the variant allele: *m_mut_* = *m_mut_* /(*m_mut_* + *UM_mut_*). The wild type allele methylation rate, *m_WT_*, is confounded by signal from normal contaminating cells and must be deconvolved. For this, we use a modified version of the CAMDAC *equations 2* and *3*, where the tumor methylation rate and copy number are expressed in terms of the mutant and wild type alleles.

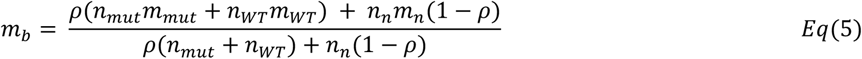

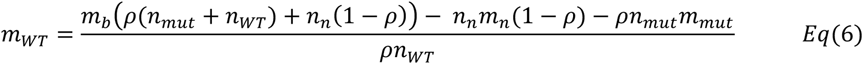

We validate CAMDAC *m_t_* by comparison with methylation estimates phased to clonal SNVs present on all copies in regions with loss of heterozygosity across our cohort (*n_mut_* = *n*_%_). At these sites, all reads reporting the variant allele can directly be assigned to the tumor cells, and methylation rates obtained from this subset of reads should be an unbiased estimate of the purified tumor methylation rate (i.e. *n_mut_* = 0). Overall, 4,485 CpG loci met these criteria. A high correlation was observed between these SNV deconvoluted *m_t_* values and CAMDAC estimates (Pearson correlation = 0.97).

### Tumor-normal differential methylation analysis

We designed a statistical test to robustly identify DMPs between tumor and normal. The number of methylated reads at a CpG is often described as a Beta-Binomial distribution where the mean of the probability distribution, the methylation rate, follows a Beta distribution. We can therefore model the observed bulk tumor and matched normal methylation rate at CpG locus i as follows:

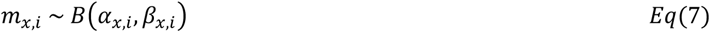

where: *m_x,i_* is the methylation rate at the ith CpG locus

*α_x,i_* is the counts methylated at the ith CpG locus

*β_x,i_* is the counts unmethylated at the ith CpG locus

*x* is *b* for the bulk tumor, *n* for the normal and *t* for the purified tumor

Our statistical design aims to test whether or not the tumor methylation rate at the ith CpG locus, *m_t,i_*, is different from the normal methylation rate, *m_n,i_*. This is depicted in equations *8* and *9*.

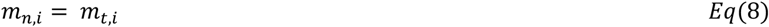

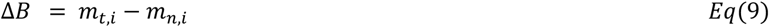

We recall from *Eq(2)* that the tumor methylation rate, *m_t_*, can be expressed as a function of *m_b_* and *m_t_*. By substituting *Eq(2)* into *Eq(9)*, we obtain:

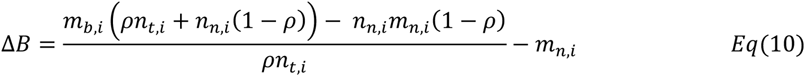

Which can be simplified to:

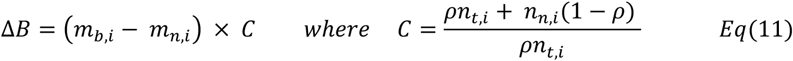

Expression (11) demonstrates that our statistical model is independent of copy number and tumor purity, which are constant for the i^th^ CpG site. However, the power to call DMPs will intrinsically depend on purity, local copy number and CpG coverage. The former two will affect the magnitude of the difference between *m_b_* and *m_t_* whilst the latter will affect the width of each beta distribution. Since there is a closed-form solution for testing *P*(*m*_*b,i*_ > *m*_*n,i*_), but not for *P*(*m*_*b,i*_ = *m*_*n,i*_) (Robinson, 2017), the null and alternate hypotheses are written as follows:

*H*_0_: *m*_*b,i*_ = *m*_*n,i*_ The methylation rate of the i^th^ CpG is identical in normal and tumor

*H*_1_: *m*_*b,i*_ > *m*_*n,i*_ The tumor is hypermethylated at this locus

*H*_2_: *m*_*b,i*_ < *m_n,i_* The tumor is hypomethylated at this locus

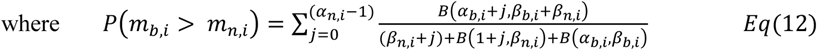

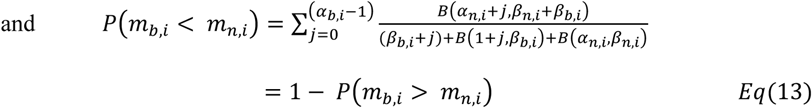

*Eq(12)* can be rewritten as follows, incorporating our *Beta*(0.5, 0.5) prior to each methylation count variable. The prior informs on the underlying methylation rate distribution and ensures finite logarithms:

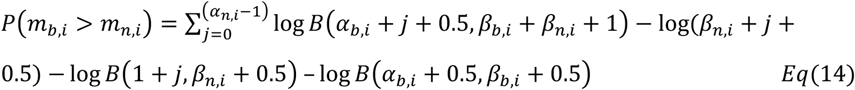

We compute the probability that a CpG site is hypo- or hypermethylated and for easier interpretation, we express these probabilities as their complement (*C* = 1-*P*), which is a measure the overlap between *m*_*b,i*_ and *m*_*n,i*_. If *C*(*m*_*b,i*_ > *m*_*n,i*_) or *C*(*m*_*b,i*_ < *m*_*n,i*_) ≤ 0.01 we accept *H_1_* or *H_2_* respectively.

Differential methylation analyses incorporating a beta distribution model reportedly show higher true positive and lower false discovery rate are obtained compared with Fisher’s and z-score methods (Raineri et al., 2014). Nevertheless, given high enough coverage, even a small difference in methylation can become statistically significant. In order to focus our analysis of biologically significant methylation changes, we require a minimum effect size of 0.2 between the purified tumor methylation rate, *m_t,i_*, and the matched normal, *m_n,i_*, for DMP calling. In theory, this allows for subclonal or allele specific changes to be picked up whilst removing spurious signal. We obtained a second set of DMPs by applying the minimum effect size threshold on the difference between the bulk tumor methylation rate, *m*_*b,i*_, and the normal methylation rate, *m*_*n,i*_ as is customary (Dolzhenko and Smith, 2014; Hansen et al., 2012; Klein and Hebestreit, 2016; Robinson et al., 2014; Wu et al., 2015), and compared the output with CAMDAC calls. The threshold of 0.2 was deemed sufficiently low to capture most mono-allelic aberrations whilst be sufficiently high to remove false positives and filter noise from the heterogeneous normal contaminating cells.

### Validation of CAMDAC purified tumor methylation rates at tumor-normal DMPs

For a given tumor sample, we validate our model by comparing observed and expected clonal bi-allelic DMPs modal methylation peak position. Beta regression was used to estimate the mode of the peak generated by hyper- and hypomethylated bi-allelic DMP populations in the bulk tumor (*m_b_*) and CAMDAC pure tumor methylation rates distribution, stratified by tumor purity, copy number and matched normal methylation status. Tumor-normal hyper- and hypomethylated CpGs were included if confidently unmethylated (99% HDI ⊆ [0, 0.2]) and methylated (99% HDI ⊆ [0.8, 1]) in the adjacent patient-matched normal, respectively. Note that allele-specific copy number states with ≤ 10,000 loci in a given sample were ignored (inter-quartile range of the number of CpGs per allele-specific copy number state is [180 125, 807 774]). CpGs on sex chromosomes were also excluded due to coverage biases against the inactive chromosome X copy shifting methylation estimates. CpGs meeting these criteria across samples had total tumor copy number ranging from 1 to 8.

The expected values of hypo- (*m*_0_) and hypermethylated (*m*_+_) clonal bi-allelic DMPs were derived from the modal methylation rate of the peaks at 0 and 1 in the patient-matched normal as estimated by beta regression as opposed to using exactly 0 and 1. The predicted CAMDAC *m_t_* are set to exactly *m*_0_ and *m*_1_ for CpGs having lost and gained methylation, respectively. The expected bulk tumor modal methylation rates at hypermethylated loci was computed by feeding sample purity, tumor copy number, *n*_n_ = 2, *m_n_* = *m*_0_ and *m_t_* = *m*_1_ into *Eq(2)*. Vice versa, *m_n_* and *m_t_* were substituted with *m*_1_ and *m*_0_, respectively, to calculate the expectation at hypomethylated loci.

### Tumor-tumor differential methylation analysis

The purified tumor methylation rate is not a beta distribution but rather the difference between two betas, the bulk tumor and normal methylation rate, scaled for tumor purity and copy number. As such, there is no exact solution to compute the highest density interval for *m_t,i_*. To address this, we simulate a credible 99% HDI for *m_t,i_* at every CpG. We use the tumor purity and CpG copy number and simulate 2000 data points for the bulk tumor (*m*_*b,i*_) and matched normal methylation rate (*m_n,i_*), given *m*_*n,i*_ ∼ *B*(*α*_*x,i*_, *β*_x,i_). Substituting these into *Eq(2)*, we obtain a vector of values for *m_t,i_* and readily extract the 99% HDI. If the purified tumor methylation rate HDI does not overlap between any two tumor regions at a given CpG and the minimum effect size, 0.2, is reached, a tumor-tumor DMP is identified.

### Tumor-normal and tumor-tumor differential methylation rate simulation framework

To appreciate the effect of tumor purity on both tumor-normal and tumor-tumor differential methylation analysis, we extracted the 20 lowest and highest purity samples in our cohort, *ρ* < 0.3 and *ρ* > 0.58, respectively. We combined the methylation information at all overlapping autosomal CpGs from these samples including the patient-matched normal of selected samples. The normal methylation rate at confidently unmethylated and methylated CpGs was extracted. Confidently unmethylated and methylated CpGs are respectively defined as having their methylation rate 99% highest density interval (HDI) boundaries in the 0.0-0.2 and 0.8-1.0 intervals (**Figure S1A-B**). This vector of values incorporates information for both distributions such as mean methylation rate and deviation as well as their respective contribution to the overall bimodal normal methylation rate profile. Sampling from this vector will yield our simulation priors.

Next, we randomly selected intra- and inter-tumor CpG pairs from samples within or across purity categories and of equal or differing total copy numbers (1, 2, 3, 4, 5 or ≥6). For each 168 possible combination of these simulation parameters, we sampled 10,000 loci from each tumor samples randomly assigned as *x* and *y* as well as their matched normal.

For each locus, we begin by obtaining the coverage information from *i*^th^ selected CpGs in sample *s* = *x* or *y*. We know the bulk tumor coverage (*cov*_*b,s,i*_), the tumor copy number (*n*_*t,s,i*_), the normal copy number (*n*_*n,s,i*_ = 2) and the global tumor purity (*ρ_s_*) and so the tumor DNA fraction (*f*_*t,s,i*_) is 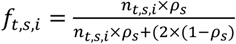. We can work out the purified tumor coverage as *cov*_*t,s,i*_ = *cov*_*b,s,i*_ × *f*_*t,s,i*_ where *cov*_*t,s,i*_ ∼ *Binom*(*cov*_b,s,i_, *f*_*t,s,i*_). The matched normal coverage (*cov*_*n,s,i*_) is therefore *cov*_*n,s,i*_ = *cov*_*b,s,i*_ − *cov*_*t,s,i*_.

We then sample a normal methylation rate prior (*p*_&,6,7_) from the confidently unmethylated (*v*_*&’(%)_) and methylated CpGs (*v*_*meth*_) from the matched normal data which is used to simulate the normal methylation rates of normal contaminating cells from both tumor samples *x* and *y*. For each sample, we obtain the counts methylated (*M_n,s,i_*) and unmethylated (*UM_n,s,i_*): *M*_*n,s,i*_ ∼ *Binom*(*cov_t,s,i_*, *p_t,s,i_*) and *UM*_*n,s,i*_ = *cov_t,s,i_* − *M*_*n,s,i*_.

The purified tumor methylation rate is obtained by randomly selecting from the same vector as the matched normal ({*p*_*n,s,i*_, *p*_*t,s,i*_} ∈ *v*_unmeth_ or {*p*_*n,s,i*_, *p*_*t,s,i*_} ∈ *v_meth_*) or from opposite vector states (*p*_*t,s,i*_ ∈ *v_unmeth_* and *p_n,i_* ∈ *v_meth_* or *p*_*t,s,i*_ ∈ *v_meth_* and *p*_n,i_ ∈ *v*_*unmeth*_). We obtain the counts methylated (*M*_*t,s,i*_) and unmethylated (*UM*_*t,s,i*_) as *M*_*t,s,i*_ ∼ *Binom*(*cov*_*t,s,i*_, *p_n,s,i_*) and *UM*_*t,s,i*_ = *cov*_*t,s,i*_ − *M*_*t,s,i*_. The bulk methylation counts for both samples are easily calculated by adding the tumor and normal counts: *M*_*bs,i*_ = *M*_*n,s,i*_ + *M*_*t,s,i*_ and *UM*_*b,s,i*_ = *UM*_*n,s,i*_ + *UM*_*t,s,i*_. In balanced copy number regions, we also simulate allele-specific DMPs whereby one allele is in the normal ground state and the other is differentially methylated. We obtain the counts methylated from the minor allele, allele A, and for the major allele, allele B, and combine them to obtain total counts methylated and unmethylated.

The expected absolute tumor-normal methylation difference at simulated bi-allelic DMPs is |*m_t_* − *m_t_*| ≈ 0.95 (**Figure 5A**). In the bulk, the magnitude of the difference depends on sample tumor purity and CpG copy number. The purified methylation rate at mono-allelic DMPs usually depends on the copy number of the mutated allele, however, in balanced copy number regions, where *n_A_* = *n_B_* and given that one copy is clonally differentially methylated and the other is in ground, 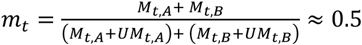. The expected tumor-normal difference at simulated mono-allelic DMPs is thus |*m_t_* − *m_t_*| ≈ 0.5 (**Figure S10A**).

Tumor-normal and tumor-tumor differential methylation calls were made using CAMDAC differential methylation analysis and the output compared with the ground truth. False negative and positive rates are obtained for the bulk and deconvolved tumor simulated data (**Figures 5B-D and S7B-D**).

### Tumor-normal and tumor-tumor differential methylation analysis on real data

We compare tumor-normal DMP calls based on CAMDAC- and FACS-purified tumor methylation rates and the bulk tumor-adjacent normal. We compute the overlap between both DMP calls, taking the ratio of DMPs called by both approaches divided by total number of FACS-purified DMPs. The overlap was 77.3% across combining DMPs across samples and the per sample overlap is displayed in (**Figure 5E**). Note that for tumor-normal DMP calling of FACS-purified data, we replace bulk tumor methylation counts, *M*_*b*_ and *UM*_*b*_ for *M*_*f,t*_ and *UM*_*f,t*_ in *Eq(14)* and apply the minimum effect size on |*m*_*f,t*_ − *m_t_*|. DMP calling power is measured as the number of tumor reads per chromosome copy (nrpcc, Dentro et al., 2021) in the bulk RRBS data, 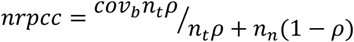.

To evaluate the impact of CAMDAC bulk tumor methylation deconvolution, we compared deconvolved with bulk tumor-tumor DMP calls between all 21 sample pairs from CRUK0062, taken for having the most tumor regions. Both DMP call sets were obtained by calculating the exact overlap between the 99% HDI of the methylation rate at the i^th^ CpG (*m*_*b,i*_) between sample pairs and applying a minimum effect size of 0.2.

To appreciate the effect of tumor purity on both inter- and intra-tumor-tumor differential methylation, we extracted the bulk and CAMDAC deconvolved tumor methylation profiles from the 20 lowest and highest purity samples in our cohort, *ρ* < 0.3 and *ρ* > 0.58, respectively. We randomly selected 1 million CpG loci from samples with the same or different purity categorical assignments both within and between patients and obtained tumor-tumor DMP calls from *m_b_* and CAMDAC *m_t_*.

### Quantifying allele-specific methylation

For all 37 normal samples in our cohort, we obtained allele-specific normal methylation rates where possible by phasing to nearby heterozygous SNPs. The normal methylation rate for the reference allele, *m*_*n,ref*_, is computed from the methylated (*M*_*n,ref*_) and unmethylated (*UM*_*n,ref*_) reads phased to the reference allele: *m*_*n,ref*_ = *M*_*n,ref*_/(*M_n,ref_* + *UM*_*n,ref*_). *Vice versa*, the alternate allele normal methylation rate, *m*_*n,alt*_, is calculated as: *m*_*n,alt*_ = *M*_*n,alt*_/(*M*_*n,alt*_ + *UM*_*n,alt*_). We then compute the exact beta posterior HDI^99^ for each allele.

Next, we assess allele-specific methylation in tumor samples. As above, we calculate bulk tumor methylation rates at the reference and alternate alleles, *m*_*b,ref*_ and *m_b,alt_*, respectively, directly from phased reads counts (un)methylated. We assign ASCAT.m clonal allele-specific copy numbers to each allele based on the BAF of the heterozygous locus used for phasing. Where multi-samples BAF phasing information is available, the segmented BAF value of the corresponding haplotype is used. If the BAF < 0.5, then major allele, *n_A_*, is the reference allele copy number, *n*_*ref*_ = *n_A_* and the minor allele, *n_B_*, is the alternate allele, *n_alt_* = *n_B_*. If BAF > 0.5, *n_alt_* = *n_A_* and *n*_*ref*_ = *n_B_*. Both the reference and alternate allele methylation rates are confounded by signal from normal contaminating cells and must be deconvolved. We modified CAMDAC *equations 2* and *3* for this purpose, where *x* is either the reference or alternate allele:

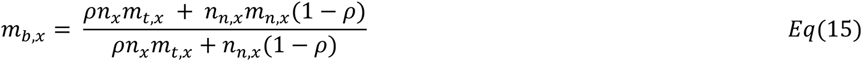

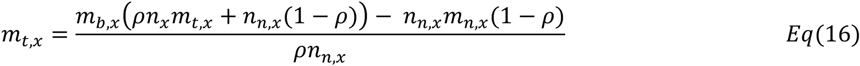

### Genomic imprinting in normal samples

We collated allele-specific methylation rate estimates at confirmed human imprinting loci from the Geneimprint database (Status=“Imprinted”, https://www.geneimprint.com). In total, 106 imprinted gene loci (2.9 genes per patient, 11 unique genes) had at least 3 non-polymorphic CpGs phased to one or more heterozygous SNPs. On average, 8.4 CpGs were close enough to be phased. We tested these CpGs for allele-specific methylation signal, requiring at least 2 consecutive CpGs with non-overlapping phased methylation rate HDI^99^ (i.e. 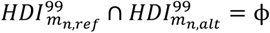) and a minimum methylation rate difference between alleles greater than 0.5 (i.e. |*m_n,ref_* − *m*_*n,alt*_| > 0.5). Overall, we detect allele-specific methylation at 100 out of the 106 loci with phased methylation information (94.3%).

### Somatic alterations at known imprinted loci

We then evaluate allele-specific pure tumor methylation at those 100 normal imprinted loci, requiring phased tumor methylation information for 3 or more non-polymorphic CpGs for inclusion.

As above, we classify tumor samples has having retained normal imprinting when at least 2 consecutive CpGs had |*m*_*t,ref*_ − *m*_*t,alt*_ | > 0.5. Tumor samples exhibiting loss of imprinting were divided into cases with and without loss of heterozygosity based on ASCAT.m allele-specific copy numbers and then further divided based on the methylation rate of the remaining allele(s). Methylated genes have mean methylation rates such that 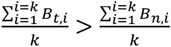 where the average phased tumor methylation rate, *B*_*t,i*_, is set to *B*_*t,i*_ = *m*_*t,x,i*_ where *x* is the remaining allele in cases with loss of heterozygosity and 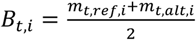 in cases without. The average allele-specific normal methylation rate at imprinted loci, *B_n,i_*, is set to 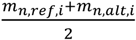. Vice versa, loci with loss of methylation have 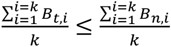.

Where distinct spatially separated genomic bins had phasing information for the same imprinted gene across tumor samples from the same patient, the most altered bin across tumor samples from the same patient was selected. For five gene loci, all but one tumor sample had phased methylation information at ≥ 3 CpGs. The imprinting status of the latter was annotated using at least one phaseable CpG and assignments were manually verified. Assignments were obtained for a total of 276 loci across genes and samples (85 genes across patients, 2.4 genes per patient, 10 unique genes).

Overall, 100/276 loci retained allele-specific methylation in tumor samples (43 loci across patients, 25/43 clonal). Loss of heterozygosity resulted in gain of methylation at 43 loci (14 loci across patients, 9/14 clonal) and demethylation at 31 loci (13 loci across patients, 8/13 clonal). Loss of imprinting without loss of heterozygosity was observed at 56 methylated loci (21 loci across patients, 10/21 clonal) and 46 unmethylated loci (20 loci across patients, 10/20 clonal). These data are summarized in a gene-wise manner for 5/10 normal imprinted genes, chosen for having phased methylation information in at least 10 tumor samples.

### Allele-specific differentially methylated CpG

Next, we obtain allele-specific pure tumor methylation rates for all CpG loci with reads overlapping with a heterozygous SNP, subset to CpGs that are confidently unmethylated in the matched normal. We compute 99% methylation rate HDI for each the reference and alternate alleles. We also evaluate the power to call an allele-specific hypermethylation on either allele for each locus given coverage, copy number and an unmethylated normal state (sampled from confidently unmethylated *m_t_* values, 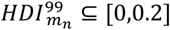). CpGs were deemed allele-specifically methylated when (1) there was sufficient power at the loci, (2) the methylation rate 99% HDIs for each allele did not overlap and (3) the between allele methylation rate difference was greater than 0.25.

For a given patient, we selected loci that were hypermethylated based on *m_t_* and/or allele-specifically methylated based on *m_t,x_* in one or more sample and powered in at least one other sample. Copy number states with less than 100 CpGs were filtered out to reduce noise. We then performed UMAP based on a matrix of the allele-specific methylation rates at these loci (R library, https://github.com/tkonopka/umap). We required at least two tumor sample per patient and a minimum of 40 CpGs for clustering (32 / 38 patients pass this criteria). The UMAP output was fed through hierarchical clustering to identify epimutation clusters (n=20), and then clusters were merged based on mean *m_t_* and *m_t,x_* values. Clusters with mean between allele difference *m*_*t*,1_ − *m*_*t*,1_< 0.125 were classed as bi-allelic. Of these, groups with *m_t_* > 0.85 ± 0.125 were classed as clonal bi-allelic. We refer to the mean methylation rate of clonal bi-allelic alterations as 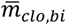. Of the remaining, only those where one allele was fully unmethylated (*m_t,i_* < 0.125) and the other was methylated on at least one copy (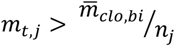, where *n_j_* is the allele specific copy number of allele *j*) were classed as clonally allele-specifically methylated. Those where less than one copy was methylated were classed as subclonal allele-specific epimutations (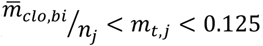). Epimutations where both alleles were partially and differentially methylated were unclassified and warrant further investigation. These may represent dynamic loci with variable methylation.

Excluding unclassified clusters, we readily obtain the relative number of (sub)clonal mono- and bi-allelic epimutations. The allele-specificity ratio is defined as the fraction of alterations that are mono-allelic and the clonality or intra-tumor heterogeneity score is the fraction of DMPs that are clonal. We fit a binomial generalized linear model (glm v4.0.5) to each of these 3 dependent variables: the fractions of allele-specific, clonal and clonal bi-allelic mutations based on histological subtype, tumor stage (IA and IB = “early”, “late” for all later stages), recurrence, SBS4 exposure (exposure > 0.35 = “high”, “low” otherwise), WGD status (majority vote) and the number of sequenced tumor samples (n ≤ 3 samples : “high”, low otherwise). We get the odds ratio for each covariate and the 95% confidence interval.

### Identifying tumor-normal differential methylated regions

First, CpGs are grouped into bins. CpGs which fall within 100bp of one another are grouped together. For each bin, we compute the number of consecutive DMPs with effect size 0.2 and p < 0.01 and the total number of DMPs. Genomic bins with 4 or more consecutive DMPs and at least 5 DMPs in total are deemed differentially methylated. Each bin is annotated for downstream analyses. Annotated gene features include CpG Islands, shores and shelves, exons, introns, 5’UTR, 3’UTR, promoters and enhancers. CpG islands, intragenic features and enhancers genomic coordinates are pulled from Ensembl via biomaRt. CpG island shores and shelves are respectively defined as the regions 0-2kb and 2kb-5kb either side of a CpG island. Intragenic annotations are simplified for each gene to the GENCODE basic transcript set. Each gene transcript promoter is defined as starting 2.5kb upstream and ending 250bp downstream of the transcription start site. For this work, we used CAMDAC with hg19 annotation set, but hg38 is also available. Note that any given CpG cluster can be associated to several features.

To validate our DMR calls, we leverage SNV purified tumor methylation rates. First, we perform tumor-normal DMP calling following the same logic as in *Eq*(14), but this time computing the probability that *P*(*m_mut_* > *m_t_*), *P*(*m_WT_* > *m_t_*) and their complements. Due to the reduced power but increased accuracy of phased methylation estimates, we decreased the threshold for DMR calling to at least 2 consecutive DMPs and obtained tumor-normal DMP calls for each allele. To confirm that DMRs that were only detected on one allele were indeed allele-specific and not due to lack of power on one allele, we identified allele-specific differentially methylated regions (AS-DMRs). CpGs were deemed allele-specifically methylated when (1) there was sufficient power at the loci, (2) the methylation rate 99% HDIs for each allele did not overlap and (3) the between allele methylation rate difference was greater than 0.2. Genomic regions with 2 or more consecutive non-polymorphic CpGs with allele-specific methylation were deemed allele-specific. Combining per allele tumor-normal DMR and AS-DMR calls, we then classified DMRs. Bins which were not AS-DMR and which where differentially methylated on both the mutant and wild type alleles were confirmed to be present on both alleles where both alleles, only the mutant allele (*in-cis*) or only the wild type allele (*in-trans*). DMRs in regions with loss of the wild type allele (*n*_EF_ < 0.5) are categorized separately as it is not possible to determine whether the DMR was *in-cis, in-trans* or on both alleles prior to the loss.

### Dimensional reduction of CAMDAC pure tumor methylation profiles

We compiled a list of all CpGs which fell within a promoter-associated tumor-normal DMR in at least one sample based on CAMDAC purified methylomes (totaling 8,570 gene promoters) and obtained mean methylation rates across CpGs for each of those genomic regions for each sample in this cohort (122 tumor and 37 normal lung samples). We applied UMAP dimensionality reduction with the R package umap (https://github.com/tkonopka/ umap) to the resulting promoter methylation profile of each sample. We map histologies (LUAD, LUSC, normal), sex and patient IDs to UMAP coordinates. We repeated this analysis selecting promoter regions based on the bulk tumor-normal DMR calls (totaling 8,387 gene promoters) feeding the mean bulk tumor methylation rates into the UMAP.

### Phylogenetic analysis

To construct phylogenetic trees for each patient, we first selected CpGs hypermethylated in at least one sampled tumor region and binned methylation rates: “0” for rate ≤ 0.3 to indicate unmethylated, “1” for rate ≥ 0.6 to indicate hypermethylated and the ambiguous character “-” in all other cases to represent intermediate methylation. Additionally, we applied a CCF threshold to previously published SNV data for each patient (Jamal-Hanjani et al. 2017): “0” for CCF ≤ 0.2, “1” for CCF > 0.2. Neighbor-joining trees were constructed based on binarized bulk and CAMDAC pure tumor methylation rates at DMPs and for SNVs using the pairwise hamming distance between samples with ape v5.4.1 (Paradis et al., 2004). All trees were outgroup-rooted on the normal sample.

We quantified the concordance between DMP and SNV trees using a score based on ancestral methylation states **(Figure S12A)**. Unlike common topology measures such as the Robinson-Foulds distance, this score incorporates (epi)mutational events. A perfect topology match between the SNV and DMP trees grants a score of 0, whereas scores increase above 0 with greater inconsistencies between ancestral methylation states given by the SNV tree and DMP tree topologies **(Figure S12B)**. To determine the significance of our observed DMP-SNV tree fit, we computed p-values by scoring each DMP tree using all possible branched rooted topologies in place of the patient’s SNV tree. For CRUK0062, our permutation is a random sample of up to n=10,395 possible topologies for computational feasibility. The resulting empirical p-value is interpreted as the probability of observing a DMP tree score given all possible SNV tree topologies – except for CRUK0062 where of subset of possible topologies is sampled – under the null hypothesis that there is no relationship between DMP and SNV tree topologies. This enables comparison between scores calculated using *m_b_* and CAMDAC *m_t_*.

To compare DMP and SNV tree trunks, we calculated the relative trunk lengths for each pair taking the trunk length (from normal to the most recent common ancestor) divided by the total length from the normal to the furthest leaf. The mutational signature (Alexandrov et al., 2013) deconvolution software deconstructSigs (Rosenthal et al., 2016) was used to extract exposures for unique mutations given SBS1, SBS2, SBS4, SBS5 and SBS13 signature profiles (COSMIC v2). A minimum of 50 mutations was required for sample inclusion.

### CAMDAC with tissue-matched normal

We were unable to sequence a patient-matched tumor-adjacent normal sample for CRUK0047. Leveraging this cohort’s 37 normal lung samples, we built a healthy lung tissue reference RRBS profile using the median normal SNP coverage and methylation counts which we fed into CAMDAC alongside tumor sample CRUK0047-R2. The ploidy and purity values (*ψ_RRBS_* = 2.15, *ρ_RRBS_* = 0.40) obtained were in agreement with the matched exome data (*ψ_WES_* = 2.29, *ρ_WES_* = 0.31). UMAP performed on CAMDAC purified tumor methylomes found CRUK0047-R2 in the same cluster as other LUAD samples. We conclude that genome-wide RRBS relative SNP coverage and methylation levels in normal lung are consistent between different patients and that tissue-matched normal RRBS data is a suitable alternative reference for CAMDAC when patient-matched data is unavailable.

## TRACERX CONSORTIUM

Charles Swanton^1,2^, Mariam Jamal-Hanjani^1,7^, Nicholas McGranahan^1,9^, Allan Hackshaw^10^, Yin Wu^1,2,3,4^, Dhruva Biswas^1,2,5^, Mihaela Angelova^2^, Stefan Boeing^6^, Takahiro Karasaki^1,2^, Selvaraju Veeriah^1^, Sonya Hessey^1,7^, James Reading^1,8^, Maise Al-Bakir^2^, Sergio A. Quezada^1,8^, Nicolai J Birkbak^11^ Gillian Price^12^, Mohammed Khalil^12^, Keith Kerr^12^, Shirley Richardson^12^, Heather Cheyne^12^, Tracey Cruickshank^12^, Gareth A Wilson^13^, Rachel Rosenthal^13^, Hugo Aerts^14, 15^, Madeleine Hewish^16^, Girija Anand^17^, Sajid Khan^17^, Kelvin Lau^18^, Michael Sheaff^18^, Peter Schmid^18^, Louise Lim, John Conibear^18^, Roland Schwarz^19^, Tom L Kaufmann^19^, Matthew Huska^19^, Jacqui Shaw^20^, Joan Riley^20^, Lindsay Primrose^20^, Dean Fennell^20,21^, Allan Hackshaw^22^, Yenting Ngai^22^, Abigail Sharp^22^, Oliver Pressey^22^, Sean Smith^22^, Nicole Gower^22^, Harjot Kaur Dhanda^22^, Kitty Chan^22^, Sonal Chakraborty^22^, Kevin Litchfield^1^, Krupa Thakkar^1^, Jonathan Tugwood^23^, Alexandra Clipson^23^, Caroline Dive^23,24^, Dominic Rothwell^23,24^, Alastair Kerr^23,24^, Elaine Kilgour^23,24^, Fiona Morgan^25^, Malgorzata Kornaszewska^25^, Richard Attanoos^25^, Helen Davies^25^, Katie Baker^26^, Mathew Carter^26^, Colin R Lindsay^26^, Fabio Gomes^26^, Fiona Blackhall^24,26^, Lynsey Priest^24,26^, Matthew G Krebs^24,26^, Anshuman Chaturvedi^24,26^, Pedro Oliveira^24,26^, Zoltan Szallasi^27^, Gary Royle^28^, Catarina Veiga^28^, Marcin Skrzypski^29^, Roberto Salgado^30^, Miklos Diossy^31^, Alan Kirk^32^, Mo Asif^32^, John Butler^32^, Rocco Bilancia^32^, Nikos Kostoulas^32^, Mathew Thomas^32^, Mairead MacKenzie^33^, Maggie Wilcox^33^, Apostolos Nakas^21^, Sridhar Rathinam^21^, Rebecca Boyles^21^, Mohamad Tufail^21^, Amrita Bajaj^21^, Keng Ang^21^, Mohammed Fiyaz Chowdhry^21^, Michael Shackcloth^34^, Julius Asante-Siaw^34^, Angela Leek^35^, Nicola Totten^35^, Jack Davies Hodgkinson^35^, Peter Van Loo^36^, William Monteiro^37^, Hilary Marshal^37^, Kevin G Blyth^38^, Craig Dick^38^, Charles Fekete^15^, Eric Lim^39^, Paulo De Sousa^39^, Simon Jordan^39^, Alexandra Rice^39^, Hilgardt Raubenheimer^39^, Harshil Bhayani^39^, Morag Hamilton^39^, Lyn Ambrose^39^, Anand Devaraj^39^, Hemangi Chavan^39^, Sofina Begum^39^, Silviu I Buderi^39^, Daniel Kaniu^39^, Mpho Malima^39^, Sarah Booth^39^, Andrew G Nicholson^39^, Nadia Fernandes^39^, Pratibha Shah^39^, Chiara Proli^39^, John Gosney^40^, Sarah Danson^41^, Jonathan Bury^41^, John Edwards^41^, Jennifer Hill^41^, Sue Matthews^41^, Yota Kitsanta^41^, Jagan Rao^41^, Sara Tenconi^41^, Laura Socci^41^, Kim Suvarna^41^, Faith Kibutu^41^, Patricia Fisher^41^, Robin Young^41^, Joann Barker^41^, Fiona Taylor^41^, Kirsty Lloyd^41^, Jason Lester^42^, Mickael Escudero^43^, Aengus Stewart^43^, Andrew Rowan^43^, Jacki Goldman ^43^, Richard Kevin Stone^43^, Tamara Denner^43^, Emma Nye^43^, Maria Greco^43^, Jerome Nicod^43^, Clare Puttick^43^, Katey Enfield^43^, Emma Colliver^43^, Alastair Magness^43^, Chris Bailey^43^, Krijn Dijkstra^43^, Vittorio Barbè^43^, Roberto Vendramin^43^, Judit Kisistok^43^, Mateo Sokac^43^, Jonas Demeulemeester^43^, Elizabeth Larose Cadieux^43^, Carla Castignani^43^, Hongchang Fu^43^, Kristiana Grigoriadis^43^, Claudia Lee^43,44^, Foteini Athanasopoulou^1,43^, Crispin Hiley^1,43^, Lily Robinson^45^, Tracey Horey^45^, Peter Russell^45^, Dionysis Papadatos-Pastos^45^, Sara Lock^46^, Kayleigh Gilbert^46^, Kayalvizhi Selvaraju^47^, Paul Ashford^47^, Oriol Pich^47^, Thomas B K Watkins^47^, Sophia Ward^47^, Emilia Lim^47^, Alexander M Frankell^47^, Christopher Abbosh^47^, Robert E Hynds^47^, Mariana Werner Sunderland^47^, Karl Peggs^47^, Teresa Marafioti^47^, John A Hartley^47^, Helen Lowe^47^, Leah Ensell^47^, Victoria Spanswick^47^, Angeliki Karamani^47^, David Moore^47^, Stephan Beck^47^, Olga Chervova ^47^, Miljana Tanic^47^, Ariana Huebner^47^, Michelle Dietzen^47^, James RM Black^47^, Carlos Martinez Ruiz^47^, Robert Bentham^47^, Cristina Naceur-Lombardelli^47^, Haoran Zhai^47^, Nnennaya Kanu^47^, Francisco Gimeno-Valiente^47^, Supreet Kaur Bola^47^, Ignacio Garcia Matos^47^, Mansi Shah^47^, Felip Galvez Cancino^47^, Despoina Karagianni^47^, Maryam Razaq^47^, Mita Akther^47^, Diana Johnson^47^, Joanne Laycock^47^, Elena Hoxha^47^, Benny Chain^47^, David R Pearce^47^, Kezhong Chen^47^, Javier Herrero^47^, Fleur Monk^47^, Simone Zaccaria^47^, Neil Magno^47^, Paulina Prymas^47^, Antonia Toncheva^47^, Monica Sivakumar^47^, Olivia Lucas^47,48^, Mark S. Hill^43,47^, Othman Al-Sawaf^47^, Seng Ung (Anakin) ^48^, Sam Gamble^48^, Sophia Wong^48^, David Lawrence^48^, Martin Hayward^48^, Nikolaos Panagiotopoulos^48^, Robert George^48^, Davide Patrini^48^, Mary Falzon^48^, Elaine Borg^48^, Reena Khiroya^48^, Asia Ahmed^48^, Magali Taylor^48^, Junaid Choudhary^48^, Sam M Janes^48^, Martin Forster^48^, Tanya Ahmad^48^, Siow Ming Lee^48^, Neal Navani^48^, Marco Scarci^48^, Pat Gorman^48^, Elisa Bertoja^48^, Robert CM Stephens^48^, Emilie Martinoni Hoogenboom^48^, James W Holding^48^, Steve Bandula^48^, Ricky Thakrar^48^, James Wilson^48^, Mansi Shah^48^, Marcos^48^, Vasquez Duran^48^, Maria Litovchenko^48^, Sharon Vanloo^48^, Piotr Pawlik^48^, Kerstin Thol^48^, Babu Naidu^49^, Gerald Langman^49^, Hollie Bancroft^49^, Salma Kadiri^49^, Gary Middleton^49^, Madava Djearaman^49^, Aya Osman^49^, Helen Shackleford^49^, Akshay Patel^49^, Christian Ottensmeier^50^, Serena Chee^50^, Aiman Alzetani^50^, Judith Cave^50^, Lydia Scarlett^50^, Jennifer Richards^50^, Papawadee Ingram^50^, Emily Shaw^50^, John Le Quesne^51^, Alan Dawson^52^, Domenic Marrone^52^, Sean Dulloo^52^, Claire Wilson^52^, Yvonne Summers^53^, Raffaele Califano^53^, Rajesh Shah^53^, Piotr Krysiak^53^, Kendadai Rammohan^53^, Eustace Fontaine^53^, Richard Booton^53^, Matthew Evison^53^, Stuart Moss^53^, Juliette Novasio^53^, Leena Joseph^53^, Paul Bishop^53^, Helen Doran^53^, Felice Granato^53^, Vijay Joshi^53^, Elaine Smith^53^, Angeles Montero^53^, Phil Crosbie^53,54^

^1^Cancer Research UK Lung Cancer Centre of Excellence, University College London Cancer Institute, London, UK

^2^Cancer Evolution and Genome Instability Laboratory, The Francis Crick Institute, London, UK

^3^Peter Gorer Department of Immunobiology, School of Immunology & Microbial Sciences, King’s College London, London, UK

^4^Immunosurveillance Laboratory, The Francis Crick Institute, London, UK

^5^Bill Lyons Informatics Centre, University College London Cancer Institute, London, UK

^6^Bioinformatics & Biostatistics and Software Development & Machine Learning Team, The Francis Crick Institute, London, UK

^7^Cancer Metastasis Lab, University College London Cancer Institute, London, UK

^8^Cancer Immunology Unit, Research Department of Haematology, University College London Cancer Institute, London, UK

^9^Cancer Genome Evolution Research Group, University College London Cancer Institute, London, UK

^10^Cancer Research UK & University College London Cancer Trials Centre, University College London, London, UK

^11^Aarhus University, Denmark

^12^Aberdeen Royal Infirmary, Aberdeen, UK

^13^Achilles Therapeutics UK Limited, London UK

^14^Artificial Intelligence in Medicine (AIM) Program, Harvard Medical School, Boston, USA

^15^Radiology and Nuclear Medicine, CARIM & GROW, Maastricht University, Maastricht, The Netherlands.

^16^Ashford and St. Peter’s Hospitals NHS Foundation Trust, Chertsey, UK.

^17^Barnet & Chase Farm Hospitals, Barnet, UK

^18^Barts Health NHS Trust, London, UK

^19^Berlin Institute for Medical Systems Biology, Max Delbrück Center for Molecular Medicine in the Helmholtz Association (MDC), Berlin, Germany

^20^Cancer Research Centre, University of Leicester, Leicester, UK

^21^Leicester University Hospitals, Leicester, UK

^22^Cancer Research UK & UCL Cancer Trials Centre, London, UK

^23^Cancer Research UK Manchester Institute, University of Manchester, Manchester, UK

^24^Cancer Research UK Lung Cancer Centre of Excellence, University of Manchester, Manchester, UK

^25^Cardiff & Vale University Health Board, Cardiff, UK

^26^Christie NHS Foundation Trust, Manchester, UK

^27^Danish Cancer Society Research Centre, Denmark

^28^Department of Medical Physics and Bioengineering, University College London Cancer Institute, London, UK

^29^Department of Oncology and Radiotherapy, Medical University of Gdańsk, Gdańsk, Poland

^30^Department of Pathology, GZA-ZNA Antwerp, Belgium

^31^Department of Physics of Complex Systems, ELTE Eötvös Loránd University, Budapest, Hungary

^32^Golden Jubilee National Hospital, Clydebank, UK

^33^Independent Cancer Patients Voice, London, UK

^34^Liverpool Heart and Chest Hospital NHS Foundation Trust, Liverpool, UK

^35^Manchester Cancer Research Centre Biobank, Manchester, UK

^36^MD Anderson Cancer Center, Houston, USA

^37^National Institute for Health Research Leicester Respiratory Biomedical Research Unit, Leicester, UK

^38^NHS Greater Glasgow and Clyde, Glasgow, UK

^39^Royal Brompton and Harefield NHS Foundation Trust, London, UK

^40^Royal Liverpool University Hospital, Liverpool, UK

^41^Sheffield Teaching Hospitals NHS Foundation Trust, Sheffield, London, UK

^42^Swansea Bay University Health Board, Swansea, UK

^43^The Francis Crick Institute, London, UK

^44^University College London Medical School, London, UK

^45^The Princess Alexandra Hospital NHS Trust, Harlow, UK

^46^The Whittington Hospital NHS Trust, London, UK

^47^University College London Cancer Institute, London, UK

^48^University College London Hospitals, London, UK

^49^University Hospital Birmingham NHS Foundation Trust, Birmingham, UK

^50^University Hospital Southampton NHS Foundation Trust, Southampton, UK

^51^University of Glasgow, Institute of Cancer Sciences, Glasgow, UK

^52^University of Leicester, Leicester, UK

^53^Wythenshawe Hospital, Manchester University NHS Foundation Trust, Manchester, UK

^54^Division of Infection, Immunity and Respiratory Medicine, University of Manchester, Manchester, UK

